# New viruses of *Cladosporium* sp. expand considerably the taxonomic structure of *Gammapartitivirus* genus

**DOI:** 10.1101/2023.06.06.543874

**Authors:** Augustine Jaccard, Nathalie Dubuis, Isabelle Kellenberger, Justine Brodard, Sylvain Schnee, Katia Gindro, Olivier Schumpp

## Abstract

Despite the fact that *Cladosporium* sp. are ubiquitous fungi, their viromes have been little studied. By analysing a collection of Cladosporium fungi, two new partitiviruses named Cladosporium cladosporioides partitivirus 1 (CcPV1) and Cladosporium cladosporioides partitivirus 2 (CcPV2) co-infecting a strain of *Cladosporium cladosporioides* were identified. Their complete genome consists in two monocistronic dsRNA segments (RNA1 and RNA2) with a high percentage of pairwise identity on 5’ and 3’ end. The RNA dependant RNA polymerase (RdRp) of both viruses and the capsid protein (CP) of CcPV1 display the classic characteristics required for their assignment to the *Gammapartitivirus* genus. In contrast, CcPV2 RNA2 encodes for a 41 KDa CP that is unusually small with a low percentage of amino acid identity as compared to CPs of other viruses classified in this genus. This sequence was used to annotate fifteen similar viral sequences with unconfirmed function. The phylogeny of the CP was highly consistent with the phylogeny of their corresponding RdRp, supporting the organization of gammapartitiviruses into three distinct clades despite stretching the current demarcation criteria.

## Introduction

A large number of different microorganisms, such as filamentous fungi, yeasts, viruses or bacteria, naturally colonise the vine [1, 2]. These organisms, collectively known as the plant microbiome, develop interactions with each other and with their hosts, all contributing to the functioning and evolution of a discrete ecological entity referred to as the holobiont [3–7]. These interactions can influence plant growth, response to pathogens, metabolite productions and adaptation to environmental changes [1, 8].

The holobiont protagonists combine different levels of interaction and the presence of mycoviruses infecting endophytes may sometimes favour the development of the host plant with potentially interesting agronomical consequences. Seminal work has demonstrated the role of the mycovirus Cryphonectria hypovirus 1 in reducing the virulence of *Cryphonectria parasitica*, the fungus responsible for chestnut blight fungus [9, 10]. A more recent study showed that the mycovirus Sclerotinia sclerotiorum hypovirulence-associated DNA virus 1 down-regulates pathogenicity factors of its fungal host, *Sclerotinia sclerotiorum*, resulting in a reduction in fungal virulence and conferring it beneficial endophytic properties that stimulates plant growth and response to stress [11]. Hence, mycoviruses appear as putative solutions for plant protection and especially for vine cultivation that requires quantities of phytosanitary products with a strong impact on natural ecosystems. These mycoviruses are particularly abundant in the grapevine, where their great diversity has been revealed by high-throughput sequencing analyses [12–14].

*Cladosporium*, one of the largest genera of dematiaceous fungi present in the environment, is also dominant as grapevine endophyte [15–17]. Its presence on leaves and berries increases progressively with the growing season [18] and late harvesting can favour the development of Cladosporium rot on the berries (*C. cladosporioides* and *C. herbarum*) which affects wine quality [19]. We analysed the prevalence and genome of viruses of this ubiquitous fungal species to understand better their role and the possible exchanges of viruses within vine fungal communities.

An approach based on the extraction of virion-associated nucleic acids (VANA), originally adapted for plant viruses, has proved highly effective on fungal mycelium. Four genomic segments forming two *Partitivirus* coexisting in *Cladosporium* sp. were identified and characterized.

This work enabled us to assign a structural function to 15 hypothetical viral proteins that form a distinct clade in the genus *Gammapartitivirus*. It includes viruses with a novel capsid protein sequence showing very little similarities to the capsid protein sequences of any other Gammapartitivirus.

## Material and method

### Fungal isolates

The fungal community was isolated from sap bleedings collected on an Agroscope experimental plot at Leytron (VS, Switzerland). The sap was collected in 50mL brown glass bottles. Bottles were sterilized with ethanol, sealed with parafoil and left for 2 weeks during the bleeding season. 100µL of sap sample diluted hundred times with sterile water were plated on Potatoes Dextrose Agar with aureomycin 12 mg.L^-1^ (PDAa). Fungi were isolated by subculture of emerging mycelium on PDA petri dishes. In total, 249 fungal isolates were cultured from the grapevine bleeding sap of 41 vinestocks. A visual classification based on the morphology of the colonies revealed a predominance of *Cladosporium* and *Aureobasidium* as previously reported on grapevines [16, 18]. To avoid the analysis of individuals from the same lineage, only one isolate of each genus was selected per plant. Their identity was confirmed by ITS sequencing of a representative isolate using ITS1F/ITS4 primer as previously described [20]. Twenty-two *Aureobasidium* isolates, 14 *Cladosporium* isolates and 12 isolates representing the diversity of colony morphotypes were selected for virus screening.

Following the identification of a virus in a *Cladopsorium* strain, 12 *Cladopsorium* isolates present in Agroscope’s fungal collection (www.mycoscope.ch/) were further screened by RT-PCR for the presence of this virus.

### Semi-purification of virus particles

The particle purification was performed according to a previously-described protocol with some modifications [21]. Briefly, 15-30 g of fresh mycelium from Potatoes Dextrose Agar culture were ground into small powder using liquid nitrogen and a mixer (Solvall Omni Mixer 17150 Homogenizer). The powder was supplemented with 6 vol of extraction buffer (0.5 M Tris, pH 8.2, 5% v/v Triton, 4% v/v Polyclar AT, 0.5% w/v bentonite, 0.2% v/v β-mercaptoethanol) and stirred on ice. After 20 min of homogenisation, the suspension was filtered through a double layer of cotton cloth. About 120 ml of filtrate was centrifuged at 4,000 rpm for 20 min. The supernatant was then collected and placed on 5 ml of 20% sucrose cushion (diluted in 0.1 M Tris, pH 8.2) followed by centrifugation at 40,000 rpm using a Beckman Coulter SW32Ti rotor for 1.5 h. The resulting pellet was incubated overnight at 4°C in 1 ml of suspension buffer (0.02 M Tris, pH 7.0, 0.001 M MgCl_2_). Enrichment in viral particles was verified by electron microscopy using 3 µl of particles as previously described [22], using the Tecnai G2 Spirit microscope (FEI, Eindhoven).

### RNA extraction

Total RNA extraction from fungal field isolates sub-cultured on agar plates was carried out according to Akbergenov et al. (2006) with the following modifications: 0.5 cm^2^ square of mycelium (50-100 mg) was cut from the edge of the plate with a scalpel, and placed in a 1,5mL Eppendorf tube with three 3 mm glass beads and frozen in liquid nitrogen. The grinding was carried out by shaking the tubes in a Tissuelyser (Qiagen) for 60 seconds at 30 Hz. If necessary, the operation was repeated once after incubation in liquid nitrogen. 1 ml of extraction buffer (6.5 M Guanidine hydrochloride; 100mM tris HCL pH=8; 100 mM β-mercaptoethanol) was added to the tube and mixed. The samples were incubated at room temperature for 10 minutes and then centrifuged for 10 minutes at 12000 rpm at 4 °C. The supernatant was transferred to a 2 ml Eppendorf tube. After the addition of 0.5 mL of Trizol (Invitrogen) reagent and 0.2 mL of Chloroform, tubes were centrifuged 10 minutes at 12 000 rpm. The upper phase was transferred to a RNase-free 50 mL polypropylene Beckman Bottles, supplemented with an equivalent volume of isopropanol, and incubated on ice for 30 minutes. The tube was centrifuged for 20 minutes at 12,000 rpm at 4°C. The pellet was washed in 70% ethanol, dried at room temperature, resuspended in 30 ul H_2_O and stored at -80°C.

VANA from the Agroscope’s fungal collection isolate *C. cladosporioides* AGS-1338 grown on PDA medium were extracted according to the protocol initially adapted for plant virus described in [24, 25]. Briefly, 200 µL of semi-purified particles described above were treated with 1 µL of DNAse and 1 µL of RNAse (Euromedex) for 90 minutes at 37°C to remove non-encapsulated RNA and DNA as described previously by Maclot et al. (2021). 400 µL of lysis buffer from the RNeasy plant mini kit (Qiagen) and 60 µL of N-Laurylsarcosine sodium salt solution 30% were added and mixed and 500 µL of the solution were loaded on a QIAshredder spin column and further processed according to manufacturer’s recommendation.

### Library preparation, sequencing and bioinformatic analyses

Total RNA extracted from 22 isolates from *Aureobasidium* sp., 14 isolates from *Cladosporium* sp. and 12 isolates representing other fungal species were pooled with equal quantity and treated for DNase with the RNeasy mini prep kit (Qiagen). RNA quality was controlled with a BioAnalyzer (Agilent Technology). A final extract of approximately 2.6 µg was used for the preparation of the cDNA library. Small RNA library preparation was performed with TruSeq small RNA kit, and mRNA with TruSeq Stranded mRNA kit. cDNA for mRNA and small RNA were sequenced using an Illumina NextSeq High library preparation kit and sequenced on an Illumina NextSeq 550 System (Illumina, USA) in paired-end 2x75 nt reads by Fasteris (Genesupport, Switzerland). Raw reads were trimmed with BBDuk 37.64 plugin and assembled using SPAdes plugin in Geneious Prime 2019.0.4 [27, 28].

Synthesis of the cDNA and tagged-library preparation from VANA was performed as described by Candresse et al. (2014) using TruSeq™ DNA Nano kit. Library quality was controlled using a Bioanalyzer 2100 and sequenced at Fasteris (Genesupport, Switzerland) on Miseq nano kit version 2 (Illumina, USA) in 1x50+8+8 cycles. Reads trimming was carried out using BBDuk 38.37 plugin from Geneious Prime 2020.0.4 (Biomatters, Auckland), and *de novo* assembly was performed using parameters of the high sensitivity mode from Geneious assembler.

### Reconstruction of whole genomic sequences and annotation

Contigs were selected and annotated using blastn on a “in-house” mycovirus database including viral genus of previously described mycoviruses, prepared from refseq sequences present in NCBI (01.05.2020). Reads were mapped to reference sequences identified by blast and primers were designed on reads stacks to confirm the sequence of each contig by Sanger sequencing and reconstruct the full genomes by RACE PCR (**Table S1**). AMV reverse transcriptase (Promega, Switzerland) and GoTaq polymerase (Promega, Switzerland) were used for a one-step protocol. RT-PCR cycling conditions were 45 minutes at 48°, followed by 2 minutes at 94°C, then 35 cycles of 45 seconds at 94°C, 40 seconds at 55°C and 1,5 minutes at 72°C, ended by 10 minutes at 72 °C. A denaturing step was applied according to Asamizu et al. [29] before using the SMARTer RACE 5’/3’ Kit (5’ section only) according to manufacturer’s recommendations. Amplified RACE and RT-PCR products were cloned in pGEM-T, sequenced and assembled using Geneious assembler with highest sensitivity parameters. Reads were mapped on the assembled sequences to control the assembly. An extra stretch of 7 nucleotides (ACATGGG) detected in all RACE sequences but not described in the kit specification was removed.

The annotation of the selected contigs was verified by online blastn and blastx analysis. The presence and size of an Open Reading Frame (ORF) was predicted for each segment by ORF finder (ncbi.nlm.nih.gov/orffinder).

### Viral particles characterisation

Virus particles were concentrated with a 10-40% sucrose gradient prepared with a Buchler gradient maker (Buchler Instruments Inc., Fort Lee, NJ, USA) with 17.5 mL of 10% (v/v) and 17.5 mL of 40% (v/v) sucrose in the suspension buffer (0.02 M Tris, pH 7.0, 0.001 M MgCl_2_). One milliliter of virus particles was overlaid on the sucrose gradient and ultracentrifuged for 2.5 hours at 30,000 rpm at 4 °C using a Beckman Coulter SW32Ti rotor. After centrifugation, fractions of 1.8 mL from top to bottom were collected and numbered from #1 to #20. Groups of three fractions were pooled, diluted in 40 mL of suspension buffer and centrifuged 2.5 h at 40’000 rpm. The pellet was suspended in 200 µL suspension buffer. Fraction groups #10 to #12, #13 to #15 and #16 to #18 were visualized by TEM. Particles from fraction #13 to #15 were used to measure the diameter of particles with ImageJ [30]. The calculation of the mean, standard deviation and the mean comparison with a student t.test was performed in R.

### LC-MS/MS

The semi-purified particles were loaded onto a 12% (v/v) sodium dodecyl sulfate-polyacrylamide gel electrophoresis (SDS-PAGE) gel. A 100 kDa size marker was used for size estimation (Biorad, low range standards). After electrophoresis, the gel was stained with Coomassie brilliant blue R250. The resulting band of the expected size of the putative capsid of CcPV1 and CcPv2 were excised and subjected to mass spectrometry coupled to liquid chromatography (LC-MS/MS) analysis at the Centre for Integrative Genomics (University of Lausanne, Switzerland) for determination of protein sequence.

### Annotation and phylogenetic analysis

The protein sequences encoding for RdRp and CP identified in this work were aligned with the protein sequences of members of the family *Partitiviridae* available from ICTV website and other members of newly described zeta and epsilon genus [31, 32] (**Table S2)**. The Human picobirnavirus strain Hy005102 reference sequence was used as an outgroup of the RdRp tree. Alignment was performed using MUSCLE version 3.8.425 implemented in Geneious Prime 2020.0.4 with standard parameters [33]. The resulting alignment quality was verified manually and zones with alignment ambiguities were excluded for tree and distance matrix calculation. The phylogenetic tree was conducted with IQ-tree, using the optimised model for maximum likelihood method [34, 35]. Branching support was obtain with 1000 boostrap with the ultrafast method from IQ-Tree [36]. The phylogenetic trees were curated on iTol [37]. Research for protein domains was done with CDsearch of NCBI in the pfam database [38] and annotation of the sequence was done with ORF finder from NCBI to identify the proper coding region of the sequence.

## Results

### Virus identification

Despite the elevated number of reads and contigs generated by Illumina sequencing of pooled fungal RNA prepared from 48 isolates collected in Leytron, only one viral contig of 123 bp could be confirmed by RT-PCR. This contig was identified in the *C*.*ramotenellum* strain AGS-Cb3.2 only. A fragment of 101 bp was amplified with the primer set 10/89 and shared 93% identity with the NCBI sequence MN034127 reconstructed from a soil metagenome study annotated as a *Partitiviridae* sp. [39]. In the absence of reverse transcription, no amplification was observed confirming the viral replicative nature of the fragment.

New primers (193/1035) designed on the sequence MN034127 enabled the amplification of a longer fragment of 1218 nucleotides that lead to the complete genomic fragment reconstruction by RACE-PCR from AGS-Cb3.2. Read mapping on this complete sequence showed that only 15 reads covering 13% of the sequence mapped on this sequence with 100% identity. As low viral titer suggested by the weak read coverage of the sequence could be due to active silencing activity, a second Illumina sequencing targeting siRNAs from the same mixed extract was performed (**Fig. S1A**). Only one read of 28 bp from sRNAs between 15 and 30 nt in size was associated with this fragment by mapping with Bowtie 2.

The infected AGS-Cb3.2 strain could not be maintained.

Twelve Cladosporium isolates maintained in the Agroscope fungal isolate collection (https://mycoscope.bcis.ch/) were then screened by RT-PCR using the primer 10/89. The *C. cladosporioides* strain AGS-1338 isolated from a vine stock of Chasselas in 2010 in Perroy (VD, Switzerland) produced a band corresponding to the expected viral fragment size, and the viral sequence was confirmed with sanger sequencing of the purified band (**Fig. S2B**). The identity of the fungal isolate was confirmed by sequencing and blastn analysis of the ITS sequence.

### Mycovirus genomes reconstruction

A third Illumina sequencing was performed using the *C. cladosporioides* isolate AGS-1338 to reconstruct the complete genome of the virus previously detected in AGS-Cb3.2, especially the second genomic fragment of this virus whose affiliation to the *Partitiviridae* family required a bisegmented genome [40]. The sequencing strategy was based on a protocol initially adapted for plant virus using VANA instead of total RNA extracts [24]. From the sequencing data, four contigs were significantly longer with sizes of 1629, 1953, 1980 and 2139 bp (**Fig. S1B**). RT-PCR with primers designed on these four contigs confirmed their presence in the strain AGS-1338. The exact 5’ and 3’ ends of each fragment were determined with RACE-PCR, and the full genome sequences were confirmed by Sanger sequencing. Final sequences were smaller than those obtained by the bioinformatics analysis of Illumina reads with sizes of 1356, 1586, 1771 and 1705 nucleotides (**Fig. 1A**). These sequences were covered by 33.3%, 21 %, 12.4% and 27.4 % of the total reads obtained by Illumina sequencing, respectively (**Fig. S1**).

**Fig. 1:**
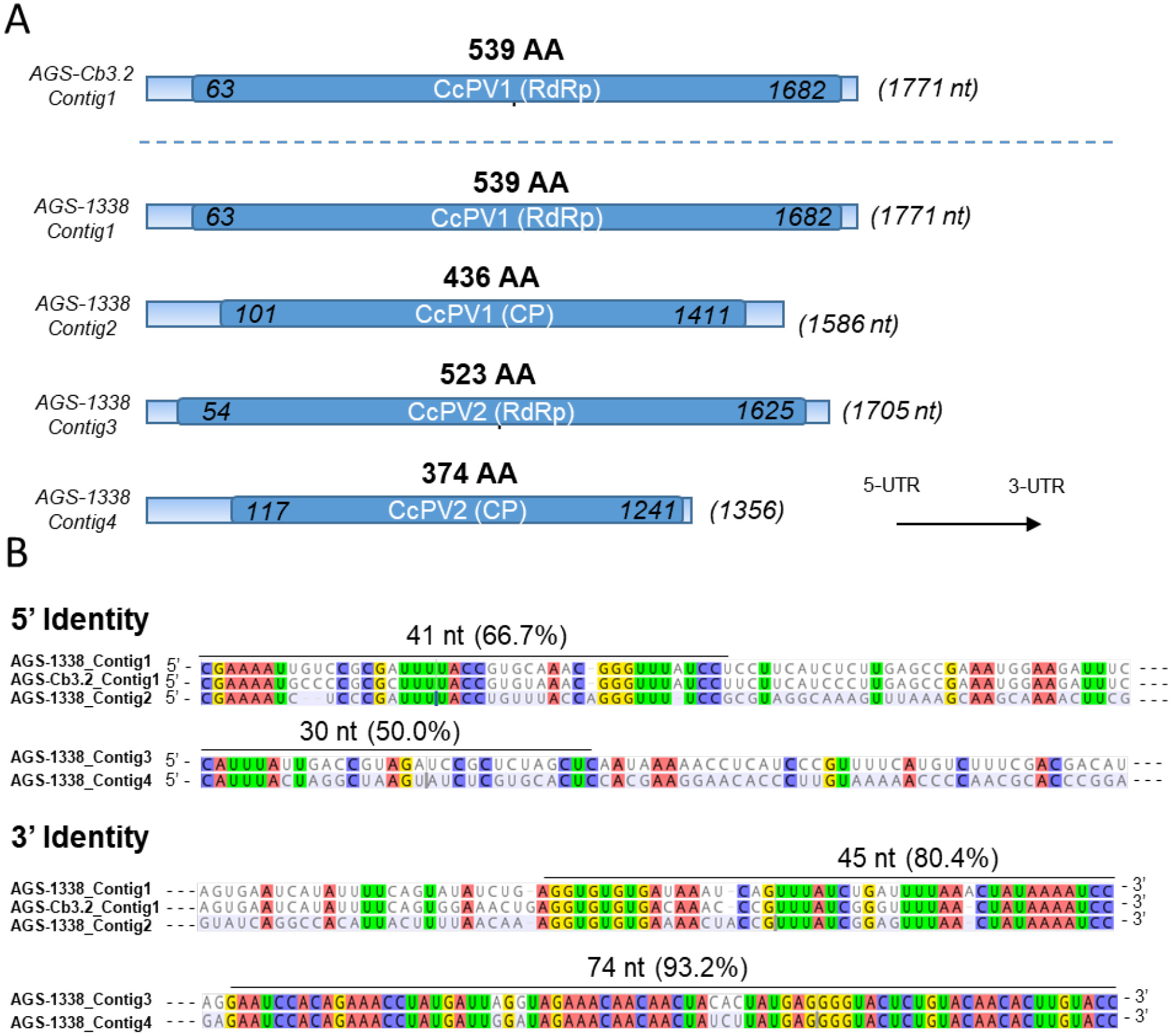
characteristic of the sequences of CcPV1 and CcPV2 detected in strain *C. cladosporioides* AGS-1338 and *C. ramotenellum* Cb3.2. A] Viral contigs. Coding sequences are highlighted in dark blue and the 5’- and 3’-UTRs sequences in light blue. Nucleic acid position of the start and end of the ORF is indicated. The size of the full nucleic sequence is in brackets. B] Alignment of the 5’- and 3’-UTRs of the genomic segments present in isolates AGS-1338 and AGS-Cb3.2. Nucleotides shared among the different sequences are highlighted and the percentage of identity of the most conserved parts is indicated.

A blastn and blastx analysis of the four contigs identified two sequences coding for RdRp (Contig1 and 3) one CP (Contig2) and a Hypothetical Protein (HP, Contig4, **Fig. 2**). The four sequences did not have a poly-A tail and showed a GC content between 45.1% and 52.7%, which corresponded to the average GC content values for dsRNA viruses in general including *Partitiviridae* [41]. An RT-like super family conserved domain was detected on Contig1 and 3 using CDsearch. The same analysis performed on the sequence reconstructed from *C. ramotenellum* strain AGS-Cb3.2 lead to the identification of a unique ORF encoding a RdRp (Contig1-Cb3.2) (**Fig. 1A**).

**Fig. 2:**
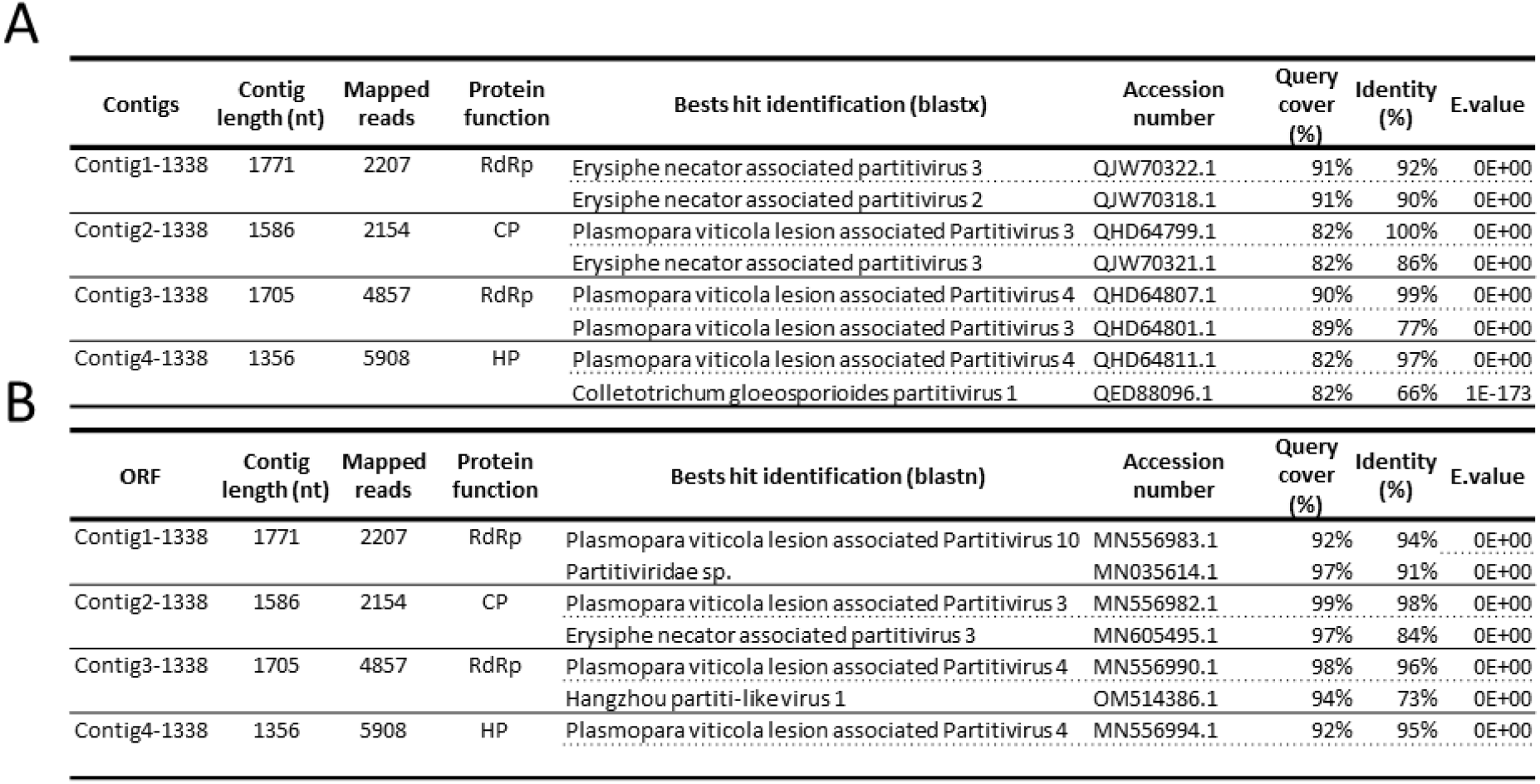
Blast annotation for the four viral segments identified in AGS-1338. A] Blastx annotation B] Blastn annotation.

All segments were more than 90% identical to one or more viral segments assigned to the *Partitiviridae* family. *Partitiviridae* are multisegmented virus, composed of two segments encoding for a RdRp (RNA1) and a CP (RNA2) [40]. This genomic organisation was confirmed for all 4 sequences by the analysis of the fragment ends. Contig1-1338 and Contig2-1338 termini showed a high sequence identity on both 5’ and 3’ ends, thereby confirming these two genomic fragments encoding for an RdRp (RNA1) and a CP (RNA2) were forming the complete genome of a virus (**Fig. 1B**). Contig1 and 2 were covered with the approximate same number of reads (**Fig. 2**).

The UTR of the Contig3-1338 and Contig4-1338 also showed high sequence identity: they shared a common stretch of 6 identical nucleotides at the 5’ end and a long stretch of 71 identical nucleotides at the 3’ (**Fig. 1B**). Contig3-1338 encoded for a RdRp. As shown by protein sequencing (see results below), the Contig4-1338 encoded for a so far undescribed CP type. Contig3 and 4 were also covered with the approximate same number of reads (**Fig. 2**). Thus, we concluded that both fragments corresponded to the RNA1 and RNA2 of a second virus infecting infecting AGS-1338. These two viruses are close to viral sequences derived from metagenomic work and are referred to as “associated” with the host species under study. In view of our work and the ubiquitous nature of Cladosporium, which is also very common on grapevines, we consider this assignment to be too uncertain. In contrast, in our work, both viruses were identified from a Cladosporium isolate in pure culture identified by sequencing and maintained in a collection. They were verified by full-length sequencing. For all these reasons, we provisionally named these viruses Cladosporium cladosporioides partitivirus 1 and 2, respectively, and hereafter refer to them as CcPV1 and CcPV2.

### Virion characterisation and CP sequencing

Particles enrichment by ultra-centrifugation was verified by TEM. Two types of particles could be observed (**Fig. 4B)**. Large dense spherical particles of 36.2±2.6 nm (n = 25) with a contrasted outline distinguished from smaller bright spherical particles of 31.5±2.4 nm (n=25) in AGS-1338 hosting CcPV1 and CcPV2. A t-test supported the size difference (p.value = 4.3e-8; **Table S3)**.

Occasionally, the concentration of CcPV1 in some plate subcultures was lower. Drawn on this finding, particle enrichments from two subcultures presenting high and low viral titre of the CcPV1 but same titre of CcPV2 were prepared (**Fig. 4**). Protein separation performed on SDS-PAGE showed two bands of about 47 and 41 kDa in the culture CcPV1^+^/CcPV2^+^ with high viral titre of both viruses, corresponding to the calculated size of the ORF from RNA 2 of CcPV1 annotated as a CP and the calculated size of the ORF from RNA 2 of CcPV2 initially annotated as HP, respectively (**Fig. 4B**). However, no band of 47 KDa was observed in the culture CcPV1^-^ /CcPV2^+^ with low viral titre of CcPV1, indicating that the missing p47 was indeed the CP of CcPV1 (**Fig. 4C**). LC-MS/MS analysis yielded 34 unique peptides covering 83% CP of CcPV1 for the p47 protein while p41 protein sequencing yielded 26 unique peptides covering 79% of the ORF of RNA 2 of CcPV2. These results demonstrated that the Contig4-1338 was a capsid protein and was therefore referred to as CP of CcPV2.

### Phylogenetic analysis

The two RdRp and CP sequences were aligned with protein sequences from representative members of the *Partitiviridae* family currently accepted by ICTV. The list was completed by viruses identified by blastx analysis of the CP for which RdRp was also available (**Table S2)**. The phylogenetic trees of both RdRps and CPs assigned CcPV1 and CcPV2 in the *Gammapartitivirus* genus. Both trees allowed the distinction of three subclades, named I, II and III, and showed a high degree of consistency for all but two viruses. Plasmopara viticola lesion associated Partitivirus 3 (PvlaPV3) had a RdRp grouped in clade III and a CP in clade II. Ustilaginoidea virens partitivirus (UvPV) is an interesting case discussed in greater detail below. Its RdRp and its CP grouped in clade II, but this virus was associated with a third protein that clustered in clade III. In both trees, CcPV1 clustered in the sub-group II and CcPV2 in clade III (**Fig. 3**).

**Fig. 3:**
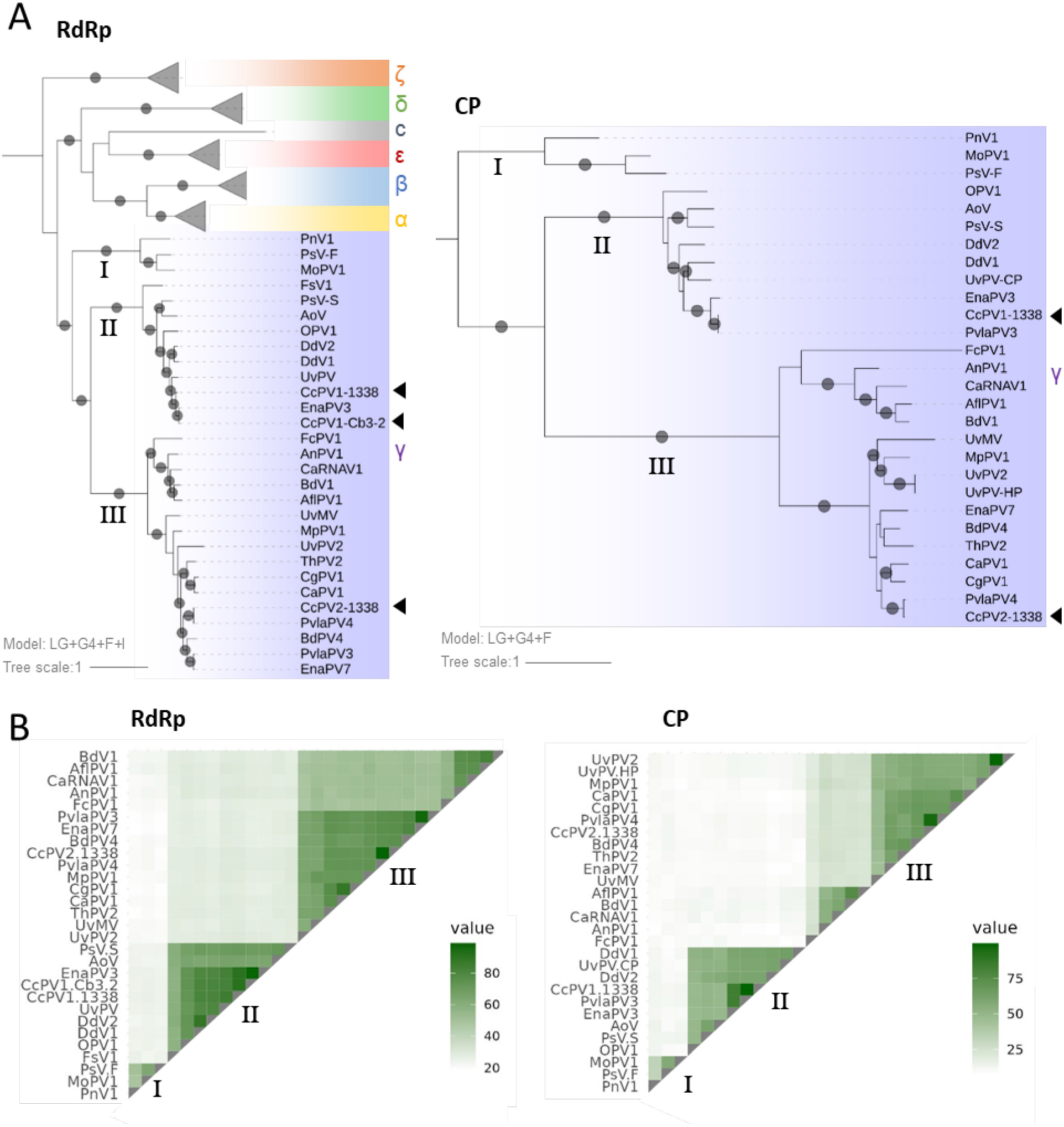
Phylogenetic tree and pairwise identity matrix of RdRp and CP proteins. A] Baesian Maximum likelywhood phylogenetic tree. Bootstrap support values greater than 70% are indicated on the branches by a grey circle. The left tree was built with LG+F+I+G4 model with a selection of RdRp sequences of Partitiviridae family (Supplementary Table S2). HBPV was used as an outgroup. The tree on the right was built with LG+G4+F model with a selection of CP sequences of the *Gammapartitivirus* family and blastx hits of CP-CcPV1 and -CcPV2. The tree was rooted according to the RdRp phylogenetic tree. B] Percentage of protein sequence identity with representative members of the *Gammapartitivirus* genus and blastx hits of CP-CcPv1 and -CcPV2 for which RdRp was also available. The numerical values of the matrix are given in Tables S4 and S5.

**Fig. 4:**
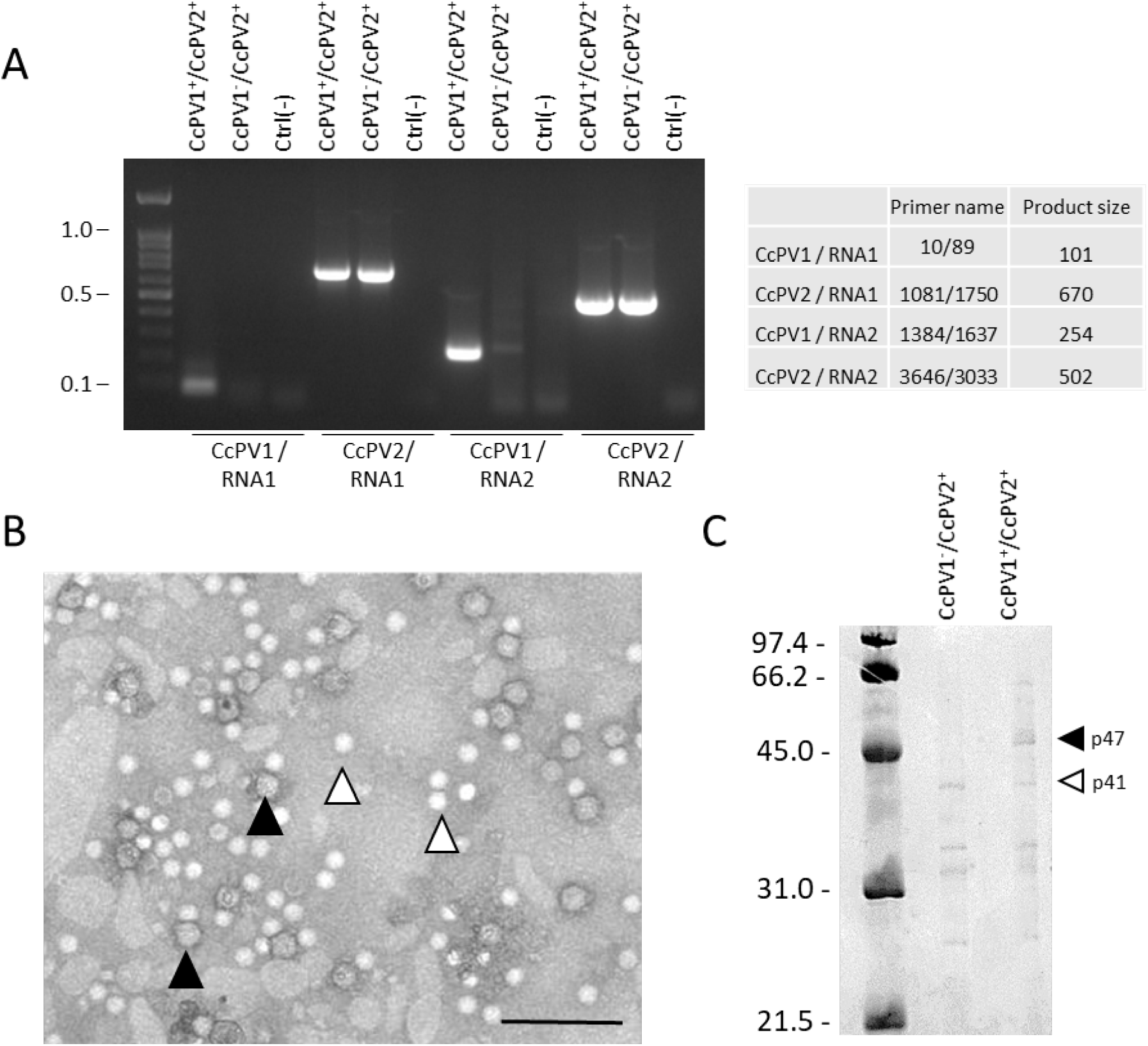
Analysis of the viral particles of *C. cladospirioides* AGS-1338. A] RT-PCR of two sub-culture of AGS-1338. DNA ladder is 100 bp. B] TEM of semi-purified viral particles from *C. cladospiroides* AGS-1338. Black arrows designate larger viral particles. White arrow designates smaller viral particles. Scale bar represents 200nm. C] SDS-PAGE of semi-purified virus particles from CcPV1^+^/CcPV2^+^ and CcPV1^-^/CcPV2^+^ subcultures. Electrophoresis gel was stained with Coomasie blue.

The RdRp identified in AGS-Cb3.2 had over 92% amino acid identity with the CcPV1 RdRp. Despite the lack of detection of a CP in AGS-Cb3.2, this high level of identity confirmed that the viral sequence detected in AGS-Cb3.2 corresponded to another isolate of CcPV1, present in another species of *Cladosporium*.

The RdRp of CcPV2 clustered in the Gamma genus with strong statistical support. However, the percentage of amino acid sequence identity fall under the 24% threshold required to delineate the *Partitiviridae* genus in pairwise comparisons with viruses of the clade I [40] (**Fig. 3B, Table S5**). The length of the RNA 2 encoding for the CP was 1356 nt that is 89 nt below the 1445 to 1611 nt limits that are currently used to delimit the genus *Gammapartitivirus*. Similarly, the length of the CP of CcPV2 (374 AA) and all viruses of group III were also well below the limits of 413 – 443 AA also used to delimit the genus *Gammapartitivirus*.

In line with these results, the calculated weight of the CP of viruses from clade III were of 40-42 KDa, close to the 41 KDa of the CP from CcPV2. This contrasted with the calculated weight of 46-48 KDa of the CP of viruses currently accepted by ICTV clustering in clade I and II. Finally, these results were also consistent with the particle sizes that could be measured: Penicillium stoloniferum viruses S (PsV-S), having particles of about 35 nm in diameter [42] was grouped in clade II with CcPV1 which had a particle size of 36.2 nm in diameter while CcPV2 having particle size of 31.5 nm in diameter was assigned to clade III.

## Discussion

High-throughput sequencing analyses revealed the presence of numerous mycoviruses representing a wide diversity of viral families in grapevine [12, 17]. Many examples of mycoviruses are capable of altering the virulence of plant pathogenic fungi [43–45] and others promote growth and/or sporulation [46–48]. However, despite some exceptions, some viral families that are very common in filamentous fungi such as the *Partitiviridae* are less frequently associated with a fungal phenotype [45, 49–51].

This apparent lack of phenotype raises questions about the role of these viruses in the development cycle of their fungal host. In order to understand better virus-fungus interactions, the aim of this study was to characterise the virome of fungal communities from grapevine wood. *Cladosporium* sp. are known to be highly represented in grapevine fungal communities and are also widespread in most ecological niches [17, 52]. This partly explains the large number of *Cladosporium* virus sequences in the NCBI databases. Nevertheless, these sequences are mainly derived from metagenomic work and form incomplete genomes in the vast majority of cases. Consequently, only seven complete viral genomes from cultivated isolates have been described to date [17, 53]. In this work, we carefully reconstructed and characterised the complete genomes of CcPV1 and CcPV2, two new mycoviruses detected in *Cladosporium* strains isolated from grapevine fungal communities. The presence of these mycoviruses was evaluated in a collection of *Cladosporium* isolates. To our knowledge, this is the first study specifically targeting *Cladosporium* isolates in pure cultures isolated from grapevine fungal communities.

Illumina sequencing of 48 pooled RNA extract prepared from petri dish cultures proved to be very insensitive: despite good RNA quality assessed by Nanodrop or Bioanalyzer and good quality data set from Illumina sequencing, only one small contig corresponding to a viral RdRp sequence of partitiviridae could be identified and confirmed by RT-PCR in a single fungal isolate (Cb3.2) of the *C. ramotenellum* species. After reconstitution of the entire genomic fragment corresponding to this contig, the read mapping produced very low coverage, reflecting a low level of expression of the mycovirus in this strain under our culture conditions. Compared to the study by Nerva et al. (2019), who detected mycoviruses in more than 15% of isolates from the grapevine fungal community using similar amounts of RNA, but prepared from liquid culture and with ribosomal depletion before library preparation, this result suggests that total RNA from 50-100 mg petri dish cultures without viral RNA enrichment is not a sufficiently sensitive method for viral genome reconstruction by high-throughput sequencing. The strain *C. ramotenellum* Cb3.2 declined rapidly and could not be maintained in collections or in liquid culture. Senescence phenomena are common in many fungal species [54, 55] and in some cases the role of a mycovirus in reducing the life span of fungal species could be demonstrated [56]. However, a stable strain of *C. cladosporioides* AGS-1338 maintained at Changins for 12 years was also infected with this mycovirus, suggesting that this viral species is not the cause of the decline of its fungal host. Thus, as for most *Partitiviridae* described so far, this mycovirus does not appear to have a negative effect on the long-term survival of its fungal host.

The sensitivity of mycovirus detection was drastically improved using a VANA enrichment from liquid culture. Four genomic segments with good coverage were identified after sequencing the VANA. None of these segments could be detected by RT-PCR in any other isolate but the RdRp of CcPV1 in isolate Cb3.2 only.

A typical and complete genomic structure of *Partitiviridae* was reconstructed for CcPV1 based on the structure and size of the genome as well as with sequence similarity with members of the genus, the ORF of RNA1-1338 (i.e. contig1) being a polymerase and the ORF of RNA2-1338 (i.e. contig2) a coat protein [57]. The ends of the 5’ and 3’ untranslated regions of the RNA1 and 2 of CcPV1 showed strong sequence identity over more than 40 nucleotides, indicating that these are two genomic segments of the same virus. The RNA1 of the CcPV1 isolate from *C. ramotenellum* (Cb3.2) and the RNA1 of the CcPV1 isolate from *C. cladosporioides* (1338) shared 40/45 (88%) and 36/42 (86%) nucleotides at the 5 and 3’ end respectively, in line with the percentage of identity of the RNA1 coding sequence (91% aa). The two genomic segments RNA1 and 2 of CcPV2 also shared highly conserved 3’ and 5’-UTR, allowing their unambiguous association (**Fig. 1**). Despite the lack of functional annotation of ORF from RNA2 resulting from blastx analysis and CD search, determination of the protein sequence of the p41 protein isolated on an acrylamide gel following purification of CcPV2 virus particles by ultracentrifugation demonstrates its role as a structural protein. This result provides experimental support for the assignment of a structural function to 15 closely related NCBI virus sequences that were previously annotated as hypothetical proteins or annotated CP with no experimental support.

This functional assignment extends the reach of this work to the taxonomic classification of the *Gammapartitivirus* genus, for which we propose to add a new clade, consisting of CcPV-2 and 15 hitherto unclassified viruses. This clade showed high levels of genetic diversity for both RdRps and CPs, down to 19% and 9%, respectively, with the most distant members of the genus. The percentage of identity for the RdRp between members of the new clade (III) and the original clade (I) is below the 24% threshold set for the genus demarcation criteria, but stands above this threshold when compared with viruses of clade II. The size of the CP is also outside the criteria that currently defines classification in the genus *Gammapartitivirus*. However, the grouping of these viruses into a new genus of *Partitiviridae* would break the existing monophyletic structure of the *Gammapartitivirus* group. Therefore, we recommend that the genus delimitation criteria for this viral family be modified to allow the incorporation of these 16 isolates representing 14 to 16 new species from the additional clade within the genus *Gammapartitivirus*. The distinction of two clades within *Gammapartitivirus* has recently been proposed, and the inclusion of 16 isolates in a third clade supports and extends these recent results by Wang et al. (2023).

The high concordance of phylogenies based on RdRp sequences on one hand and on CP sequences on the other highlights two inconsistencies for UvPV and PvlaPV3. PvlaPV3 was identified in a metagenomic study. In the absence of biologically available isolates and RACE-PCR data to compare the complete ends of the fragments, it is not possible to verify whether this inconsistency is an incorrect association of segments from two distinct viruses or cases of reassortment. UvPV is an interesting case prepared from a pure culture of a *Ustilaginoidea virens* strain maintained in a collection and containing 4 viral genomic fragments. The first two fragments associated by 5’ end analysis correspond to a group II RdRp and CP. The third fragment - unfortunately incomplete at its ends - clustered in the new clade III of gammapartitiviruses. Further work is required to verify whether these are two distinct viruses for which an RdRp is missing, or whether this third fragment is a form of virus that is a satellite of the first.

The characterisation of the CcPV2 genome also reveals an exceptionally long conserved region spanning 69/74 nucleotides of the 3’ UTR, which was almost the entire non-coding area of RdRp (84 nt) and a large part of the non-coding area of CP (118 nt). This long-conserved region of CcPV2 has no homology to any other virus. Highly conserved UTR of segments of multipartite virus have previously been observed in viruses of different families and may extend to the entire untranslated sequence [58–60]. However, in the *Partitiviridae* family, the conserved motif was so far short, restricted to a few nucleotides of the untranslated ends with some genus specificity [49, 61] although it sometimes extended beyond these few nucleotides [62, 63].

The role of high conservation level of UTRs remains to be defined, but similar to the role proposed for segmented viruses [64–66], it may ensure packaging and transcriptional specificity to limit reassortment. Thus, this high degree of specificity of the untranslated ends that distinguishes these two viral species, most likely contributes to the stability of their coexistence within the same fungal strain over the last 12 years.

## Conclusion

The two complete genomic sequences presented in this work have made it possible to extend the genus *Gammapartitivirus* by integrating a large set of previously unassigned sequences into a new clade of this viral genus. This work has also presented a new group of capsid proteins.

## Author contributions

**Augustine Jaccard:** Conceptualization, Methodology, Formal analysis, Investigation, Writing - Original Draft, Visualization

**Nathalie Dubuis:** Investigation

**Isabelle Kellenberger:** Investigation

**Justine Brodard:** Investigation

**Sylvain Schnee:** Writing - Review & Editing

**Katia Gindro:** Writing - Review & Editing

**Olivier Schumpp:** Conceptualization, Writing - Original Draft, Writing - Review & Editing, Supervision, Project administration, Funding acquisition

## Acknowledgments

We would like to thank François Maclot for his precious advices for VANA extraction. We are grateful to Nicole Lecoultre and Emilie Michellod who helped in the fungal community construction. We are also grateful to Arnaud Blouin for his careful review of the manuscript.

