## Supplementary material for "New viruses of *Cladosporium* sp. expand considerably the taxonomic structure of *Gammapartitivirus* genus": Table S

Table S1: Primer used for the viral sequence detection and reconstruction.

| Name | Sequence (5'-3') | Application | Sequence used for the design |
|---|---|---|---|
| 2 | CGCGCTTTTACCGTGTAA | RT-PCR | MN035614 |
| 10 | CCTTAGTTTGCAGCACCA | RT-PCR | Contig476912-Cb3.2 |
| 89 | CTCCTTTGATGTATCGTKACTT | RT-PCR | Contig476912-Cb3.2 |
| 193 | TCTACTTCCTGGCAGTCC | RT-PCR | MN034127 |
| 438 | GGTGGAGTGCGTCAACTA | RACE | RdRp-Cb3.2 |
| 781 | ATCTTGGTGGACGCAGAT | RT-PCR + RACE | RdRp-Cb3.2 |
| 1707 | CMCACACCTCAGTTTTCACT | RT-PCR | MN035614 |
| 1035 | GGAAACGTGGGCATGTAC | RT-PCR | MN034127 |
| 1 | ACATGGGCGAAAATTGTCC | RT-PCR | Contig1-1338 |
| 275 | AGGCTCTCTCACCTGA | RACE | Contig1-1338 |
| 736 | CGCATTGACGCTTGTTGA | RT-PCR | Contig1-1338 |
| 1349 | GGTGTGGTATTTCATAAACAC | RT-PCR | Contig1-1338 |
| 2077 | ACATGGGGGATTTTATAGTTTA | RT-PCR | Contig1-1338 |
| 18 | GGCGAAGGCATTCACTTT | RT-PCR | Contig2-1338 |
| 121 | CCTTGATCTGATCGGCGA | RACE | Contig2-1338 |
| 1087 | GCCGTCTATTCTGGAGGA | RT-PCR | Contig2-1338 |
| 1245 | ATGAGTCACACAGCTGGT | RT-PCR | Contig2-1338 |
| 1384 | TTACCAGGGTTTTCCGCG | RACE | Contig2-1338 |
| 1637 | CGGATGAAGAAGTGGCGA | RT-PCR | Contig2-1338 |
| 3033 | ATGATGTAGGAGCCCAGG | RT-PCR | Contig4-1338 |
| 3133 | GTGAAGTTGCATTCGGGA | RACE | Contig4-1338 |
| 3646 | ACTTCTCCGGTCTCGAAA | RT-PCR | Contig4-1338 |
| 2420 | CTTCTGCCTGTGCGATAA | RACE | Contig4-1338 |
| 2400 | CTTATCGCACAGGCAGAA | RT-PCR | Contig4-1338 |
| 3495 | GCCAATGATCTGAGCCAA | RACE | Contig4-1338 |
| 292 | CTTCGGTATGCGACAGAG | RT-PCR | Contig3-1338 |
| 440 | GGAGTACGAGTGTAGCCT | RACE | Contig3-1338 |
| 1081 | ATCACTGGTACCCAACGA | RT-PCR | Contig3-1338 |
| 1200 | GGTAACACAGCGACCATT | RT-PCR | Contig3-1338 |
| 1608 | TCAAGATGCGCACTTTGT | RACE | Contig3-1338 |
| 1750 | GCTTATACACGTTCGGCA | RT-PCR | Contig3-1338 |

**Table S2: Selection of viruses of the Partitiviridae family for alignment and phylogenetic tree construction.**

| Acronym | Accession (RdRp) | Accession (CP) | Accession (HP) | Virus name | Familly |
|---|---|---|---|---|---|
| HPBV | BAD98236.1 | | | Human picobirnavirus | - |
| BCV1 | YP_002308574.1 | | | Beet cryptic virus 1 | *Alphapartitivirus* |
| CCRSaPV | AJ781401 | | | Cherry chlorotic rusty spot associated partitivirus | *Alphapartitivirus* |
| CpCV1 | AM999771 | | | Chondrostereum purpureum cryptic virus 1 | *Alphapartitivirus* |
| CrCV | YP_009508046.1 | | | Carrot cryptic virus | *Alphapartitivirus* |
| FvBV | AB465308 | | | Flammulina velutipes browning virus | *Alphapartitivirus* |
| HetPV1 | HQ541323 | | | Heterobasidion partitivirus 1 | *Alphapartitivirus* |
| HetPV3 | FJ816271 | | | Heterobasidion partitivirus 3 | *Alphapartitivirus* |
| RnPV2 | YP_007419077.1 | | | Rosellinia necatrix partitivirus 2 | *Alphapartitivirus* |
| VCV | YP_272124.1 | | | Vicia cryptic virus | *Alphapartitivirus* |
| WCCV1 | AY705784 | | | White clover cryptic virus 1 | *Alphapartitivirus* |
| AhV | L39125 | | | Atkinsonella hypoxylon virus | *Betapartitivirus* |
| CCCV2 | JX971982 | | | Crimson clover cryptic virus 2 | *Betapartitivirus* |
| CCV | ET80948.1 | | | Cannabis cryptic virus | *Betapartitivirus* |
| CpPV | YP_001911122.1 | | | Ceratocystis polonica partitivirus | *Betapartitivirus* |
| CrV1 | AY603052 | | | Ceratocystis resinifera virus 1 | *Betapartitivirus* |
| DCV2 | JX971984 | | | Dill cryptic virus 2 | *Betapartitivirus* |
| FpV1 | AF047013 | | | Fusarium poae virus 1 | *Betapartitivirus* |
| HetPV2 | HM565953 | | | Heterobasidion partitivirus 2 | *Betapartitivirus* |
| HetPV7 | JN606091 | | | Heterobasidion partitivirus 7 | *Betapartitivirus* |
| HetPV8 | JX625227 | | | Heterobasidion partitivirus 8 | *Betapartitivirus* |
| HTCV2 | JX971980 | | | Hop trefoil cryptic virus 2 | *Betapartitivirus* |
| PmV1 | ABW82141.1 | | | Primula malacoides virus 1 | *Betapartitivirus* |
| PoV1 | AY533038 | | | Pleurotus ostreatus virus 1 | *Betapartitivirus* |
| RCCV2 | JX971978 | | | Red clover cryptic virus 2 | *Betapartitivirus* |
| RnV1 | YP_392480.1 | | | Rosellinia necatrix partitivirus 1-W8 | *Betapartitivirus* |
| RsV | NP_620659.1 | | | Rhizoctonia solani virus 717 | *Betapartitivirus* |
| WCCV2 | JX971976 | | | White clover cryptic virus 2 | *Betapartitivirus* |
| CSpV1 | U95995 | | | Cryptosporidium parvum virus 1 | *Cryspovirus* |
| BCV2 | YP_009508068.1 | | | Beet cryptic virus 2 | *Deltapartitivirus* |
| PCV1 | YP_009466859.1 | | | Pepper cryptic virus 1 | *Deltapartitivirus* |

| | | | | | | |
|---|---|---|---|---|---|---|
| PCV2 | YP_009351838.1 | | | Pepper cryptic virus 2 | *Deltapartitivirus* | |
| BbPV3 | QFP40245.1 | | | Beauveria bassiana partitivirus 3 | *Epsilonpartitivirus* | |
| CePV1 | AZT88590.1 | | | Colletotrichum eremochloae partitivirus 1 | *Epsilonpartitivirus* | |
| RsdRV5 | AVP26802.1 | | | Rhizoctonia solani dsRNA virus 5 | *Epsilonpartitivirus* | |
| AflPV1 | QDE53634.1 | | QDE53635.1 | Aspergillus flavus partitivirus 1 | *Gammapartitivirus* | III |
| AnPV1 | BDF97658.1 | BDF97659.1 | | Aspergillus niger partitivirus 1 | *Gammapartitivirus* | III |
| AoV | ABV30675.1 | YP_009665973.1 | | Aspergillus ochraceous virus | *Gammapartitivirus* | II |
| BdPV4 | UVZ34178.1 | UVZ34179.1 | | Botryosphaeria dothidea partitivirus 4 | *Gammapartitivirus* | III |
| BdV1 | AIE47694.1 | | AIE47695.1 | Botryosphaeria dothidea virus 1 | *Gammapartitivirus* | III |
| CaPV1 | UYD21365.1 | | UYD21364.1 | Colletotrichum associated partitivirus 1 | *Gammapartitivirus* | III |
| CaRNAV1 | AGL42312.1 | | AGL42313.1 | Colletotrichum acutatum RNA virus 1 | *Gammapartitivirus* | III |
| CcPV1 | WEU80702.1 | WEU80705.1 | | Cladosporium cladosporioides partitivirus 1 | *Gammapartitivirus* | II |
| CcPV2 | WEU80703.1 | WEU80704.1 | | Cladosporium cladosporioides partitivirus 2 | *Gammapartitivirus* | III |
| CgPV1 | QED88095.1 | | QED88096.1 | Colletotrichum gloeosporioides partitivirus 1 | *Gammapartitivirus* | III |
| CrPV1 | WEU80701.1 | - | | Cladosporium ramotenellum partitivirus 1 | *Gammapartitivirus* | III |
| DdV1 | AAG59816.1 | NP_116742.1 | | Discula destructiva virus 1 | *Gammapartitivirus* | II |
| DdV2 | AAK59379.1 | NP_620302.1 | | Discula destructiva virus 2 | *Gammapartitivirus* | II |
| EnaPV3 | QJW70322.1 | QJW70321.1 | | Erysiphe necator associated partitivirus 3 | *Gammapartitivirus* | II |
| EnaPV7 | QJW70316.1 | QJW70323.1 | | Erysiphe necator associated partitivirus 7 | *Gammapartitivirus* | III |
| FcPV1 | QOL02536.1 | | QOL02537.1 | Fusarium cerealis partitivirus 1 | *Gammapartitivirus* | III |
| FsV1 | BAA09520.1 | | | Fusarium solani virus 1 | *Gammapartitivirus* | II |
| MoPV1 | APP18151.1 | APP18152.1 | | Magnaporthe oryzae partitivirus 1 | *Gammapartitivirus* | I |
| MpPV1 | QKO02079.1 | | QKO02080.1 | Macrophomina phaseolina partitivirus 1 | *Gammapartitivirus* | III |
| OPV1 | AM087202 | YP_009508237.1 | | Ophiostoma partitivirus 1 | *Gammapartitivirus* | II |
| PnV1 | YP_009551507.1 | YP_009551508.1 | | Pythium nunn virus 1 | *Gammapartitivirus* | I |
| PsV-F | AY738336 | AAU95759.1 | | Penicillium stoloniferum virus F | *Gammapartitivirus* | I |
| PsV-S | AY156521 | YP_052857.1 | | Penicillium stoloniferum virus S | *Gammapartitivirus* | II |
| PvlaPV3 | QHD64801.1 | QHD64799.1 | | Plasmopara viticola lesion associated Partitivirus 3 | *Gammapartitivirus* | III/II |
| PvlaPV4 | QHD64807.1 | | QHD64811.1 | Plasmopara viticola lesion associated Partitivirus 4 | *Gammapartitivirus* | III |
| ThPV2 | UVB68789.1 | UVB68790.1 | | Trichoderma harzianum partitivirus 2 | *Gammapartitivirus* | III |
| UvMV | AGJ03719.1 | | AGJ03720.1 | Ustilaginoidea virens mycovirus | *Gammapartitivirus* | III |
| UvPV | AGO04402.1 | AGO4403.1 | AGO04404.1 | Ustilaginoidea virens partitivirus | *Gammapartitivirus* | II / III |
| UvPV2 | YP_008327312.1 | | YP_008327313.1 | Ustilaginoidea virens partitivirus 2 | *Gammapartitivirus* | III |

| | | | | |
|---|---|---|---|---|
| AaPV1 | APT70073.1 | Alternaria alternata partitivirus 1 | *Zetapartitivirus* | - |
| BdPV1 | AGZ84316.1 | Botryosphaeria dothidea partitivirus 1 | *Zetapartitivirus* | - |
| FePV1 | QOW77954.1 | Fusarium equiseti partitivirus 1 | *Zetapartitivirus* | - |

**Table S3: Particle diameter (nm).** Dense spherical particles with a contrasted outline and  bright spherical particles were observed in strain AGS-1338 infected and measured under TEM.

| Bright spherical particles | Dense spherical particles with a contrasted outline |
|---|---|
| 29.07 | 40.2 |
| 28.07 | 38.01 |
| 32.02 | 35 |
| 31 | 33 |
| 33.02 | 36.35 |
| 34 | 38.05 |
| 33.14 | 39 |
| 29.07 | 38.12 |
| 31 | 36 |
| 34.13 | 39 |
| 29.07 | 40.01 |
| 34.02 | 39.01 |
| 30.41 | 40.01 |
| 27.02 | 34.06 |
| 30.07 | 34 |
| 33.14 | 33.06 |
| 34.06 | 33 |
| 36.01 | 37.01 |
| 33.14 | 36.01 |
| 28.02 | 31.06 |
| 30.07 | 35 |
| 30.02 | 34.06 |
| 34.02 | 37.05 |
| 33.14 | 34 |
| | 35 |

**Table S4**: Alignment matrix of percentage of identity of CP sequence of a selection of *Gammapartitivirus*

| % aa Identity | PnV1_ | PsV-F | MoPV1 | OPV1 | PsV-S | AoV | EnaPV3 | PvlaPV3 | CcPV1-1338 | DdV2 | UvPV-CP | DdV1 | FcPV1 | AnPV1 | CaRNAV1 | BdV1 | AflPV1 | UvMV | EnaPV7 | ThPV2 | BdPV4 | CcPV2-1338 | PvlaPV4 | CgPV1 | CaPV1 | MpPV1 | UvPV-HP | UvPV2 |
|---|---|---|---|---|---|---|---|---|---|---|---|---|---|---|---|---|---|---|---|---|---|---|---|---|---|---|---|---|
| PnV1_ | | 33 | 34 | 14 | 14 | 15 | 13 | 14 | 14 | 14 | 13 | 13 | 11 | 9 | 10 | 10 | 10 | 12 | 11 | 11 | 11 | 11 | 11 | 12 | 11 | 12 | 13 | 13 |
| PsV-F | 33 | | 58 | 11 | 14 | 11 | 11 | 13 | 13 | 11 | 11 | 10 | 10 | 12 | 10 | 12 | 10 | 10 | 11 | 13 | 13 | 13 | 13 | 11 | 13 | 10 | 11 | 11 |
| MoPV1 | 34 | 58 | | 11 | 14 | 13 | 12 | 13 | 14 | 12 | 11 | 12 | 11 | 11 | 12 | 13 | 12 | 11 | 10 | 11 | 11 | 11 | 11 | 10 | 11 | 9 | 10 | 10 |
| OPV1 | 14 | 11 | 11 | | 50 | 49 | 48 | 49 | 49 | 51 | 51 | 52 | 12 | 10 | 14 | 14 | 12 | 10 | 11 | 11 | 11 | 11 | 11 | 11 | 10 | 11 | 11 | 11 |
| PsV-S | 14 | 14 | 14 | 50 | | 62 | 50 | 51 | 51 | 55 | 53 | 54 | 11 | 11 | 11 | 13 | 14 | 11 | 11 | 11 | 11 | 11 | 12 | 10 | 10 | 11 | 12 | 12 |
| AoV | 15 | 11 | 13 | 49 | 62 | | 50 | 50 | 50 | 57 | 53 | 52 | 13 | 10 | 13 | 12 | 12 | 12 | 11 | 12 | 12 | 12 | 13 | 12 | 10 | 10 | 12 | 12 |
| EnaPV3 | 13 | 11 | 12 | 48 | 50 | 50 | | 84 | 84 | 61 | 59 | 60 | 10 | 11 | 14 | 14 | 13 | 12 | 11 | 11 | 11 | 12 | 11 | 11 | 10 | 12 | 13 | 13 |
| PvlaPV3 | 14 | 13 | 13 | 49 | 51 | 50 | 84 | | 100 | 61 | 60 | 61 | 10 | 10 | 12 | 13 | 12 | 12 | 11 | 12 | 11 | 12 | 12 | 11 | 10 | 11 | 13 | 13 |
| CcPV1-1338 | 14 | 13 | 14 | 49 | 51 | 50 | 84 | 100 | | 61 | 60 | 61 | 10 | 10 | 12 | 13 | 12 | 12 | 11 | 12 | 11 | 12 | 12 | 11 | 10 | 11 | 13 | 13 |
| DdV2 | 14 | 11 | 12 | 51 | 55 | 57 | 61 | 61 | 61 | | 62 | 62 | 10 | 9 | 11 | 12 | 12 | 10 | 9 | 10 | 10 | 10 | 10 | 10 | 9 | 11 | 12 | 12 |
| UvPV-CP | 13 | 11 | 11 | 51 | 53 | 53 | 59 | 60 | 60 | 62 | | 64 | 9 | 12 | 13 | 14 | 14 | 11 | 9 | 10 | 11 | 11 | 11 | 10 | 9 | 12 | 12 | 12 |
| DdV1 | 13 | 10 | 12 | 52 | 54 | 52 | 60 | 61 | 61 | 62 | 64 | | 10 | 12 | 12 | 14 | 15 | 11 | 11 | 9 | 10 | 12 | 12 | 11 | 10 | 14 | 14 | 14 |
| FcPV1 | 11 | 10 | 11 | 12 | 11 | 13 | 10 | 10 | 10 | 10 | 9 | 10 | | 26 | 24 | 25 | 24 | 19 | 20 | 22 | 22 | 21 | 21 | 21 | 19 | 22 | 20 | 20 |
| AnPV1 | 9 | 12 | 11 | 10 | 11 | 10 | 11 | 10 | 10 | 9 | 12 | 12 | 26 | | 54 | 54 | 51 | 25 | 24 | 26 | 26 | 26 | 26 | 26 | 26 | 29 | 27 | 27 |
| CaRNAV1 | 10 | 10 | 12 | 14 | 11 | 13 | 14 | 12 | 12 | 11 | 13 | 12 | 24 | 54 | | 60 | 58 | 23 | 24 | 26 | 24 | 25 | 24 | 24 | 24 | 27 | 24 | 25 |
| BdV1 | 10 | 12 | 13 | 14 | 13 | 12 | 14 | 13 | 13 | 12 | 14 | 14 | 25 | 54 | 60 | | 76 | 25 | 23 | 24 | 24 | 25 | 25 | 25 | 24 | 26 | 26 | 26 |
| AflPV1 | 10 | 10 | 12 | 12 | 14 | 12 | 13 | 12 | 12 | 12 | 14 | 15 | 24 | 51 | 58 | 76 | | 24 | 23 | 23 | 23 | 25 | 24 | 24 | 23 | 25 | 26 | 26 |
| UvMV | 12 | 10 | 11 | 10 | 11 | 12 | 12 | 12 | 12 | 10 | 11 | 11 | 19 | 25 | 23 | 25 | 24 | | 45 | 46 | 49 | 49 | 49 | 49 | 48 | 52 | 50 | 50 |
| EnaPV7 | 11 | 11 | 10 | 11 | 11 | 11 | 11 | 11 | 11 | 9 | 9 | 11 | 20 | 24 | 24 | 23 | 23 | 45 | | 62 | 66 | 64 | 64 | 62 | 61 | 55 | 54 | 54 |
| ThPV2 | 11 | 13 | 11 | 11 | 11 | 12 | 11 | 12 | 12 | 10 | 10 | 9 | 22 | 26 | 26 | 24 | 23 | 46 | 62 | | 68 | 61 | 60 | 65 | 63 | 54 | 57 | 56 |
| BdPV4 | 11 | 13 | 11 | 11 | 11 | 12 | 11 | 11 | 11 | 10 | 11 | 10 | 22 | 26 | 24 | 24 | 23 | 49 | 66 | 68 | | 66 | 67 | 68 | 66 | 60 | 59 | 58 |
| CcPV2-1338 | 11 | 13 | 11 | 11 | 11 | 12 | 12 | 12 | 12 | 10 | 11 | 12 | 21 | 26 | 25 | 25 | 25 | 49 | 64 | 61 | 66 | | 96 | 67 | 66 | 60 | 56 | 56 |
| PvlaPV4 | 11 | 13 | 11 | 11 | 12 | 13 | 11 | 12 | 12 | 10 | 11 | 12 | 21 | 26 | 24 | 25 | 24 | 49 | 64 | 60 | 67 | 96 | | 66 | 65 | 60 | 56 | 56 |
| CgPV1 | 12 | 11 | 10 | 11 | 10 | 12 | 11 | 11 | 11 | 10 | 10 | 11 | 21 | 26 | 24 | 25 | 24 | 49 | 62 | 65 | 68 | 67 | 66 | | 73 | 58 | 56 | 56 |
| CaPV1 | 11 | 13 | 11 | 10 | 10 | 10 | 10 | 10 | 10 | 9 | 9 | 10 | 19 | 26 | 24 | 24 | 23 | 48 | 61 | 63 | 66 | 66 | 65 | 73 | | 54 | 57 | 57 |
| MpPV1 | 12 | 10 | 9 | 11 | 11 | 10 | 12 | 11 | 11 | 11 | 12 | 14 | 22 | 29 | 27 | 26 | 25 | 52 | 55 | 54 | 60 | 60 | 60 | 58 | 54 | | 61 | 61 |
| UvPV-HP | 13 | 11 | 10 | 11 | 12 | 12 | 13 | 13 | 13 | 12 | 12 | 14 | 20 | 27 | 24 | 26 | 26 | 50 | 54 | 57 | 59 | 56 | 56 | 56 | 57 | 61 | | 99 |
| UvPV2 | 13 | 11 | 10 | 11 | 12 | 12 | 13 | 13 | 13 | 12 | 12 | 14 | 20 | 27 | 25 | 26 | 26 | 50 | 54 | 56 | 58 | 56 | 56 | 56 | 57 | 61 | 99 | |

**Table S5**: Alignment matrix of percentage of identity of RdRp sequence of a selection of *Gammapartitivirus*

| % aa Identity | HPBV | HetRV2 | HetRV7 | AhV | CpPV | CrV1 | RsV | CCV | CCCV2 | PmV1 | DCV2 | HTCV2 | RCCV2 | WCCV2 | RnV1 | PoV1 | FpV1 | HetRV8 | RnPV2 | HetPV1 | CCRSaPV | BCV1 | CrCV | VCV | WCCV1 | HetRV3 | CpCV1 | FvBV | PCV1 | BCV2 | PCV2 | CSpV1 | RsdRV5 | BbPV3 | CePV1 | FePV1 | AaPV1 | BdPV1_ | PnV1 | MoPV1 | PsV-F | FsV1 | OPV1 | DdV1 | DdV2 | UvPV | CcPV1-1338 | CcPV1-Cb3-2 | EnaPV3 | AoV | PsV-S | UvPV2 | UvMV | ThPV2 | CaPV1 | CgPV1 | MpPV1 | PvlaPV4 | CcPV2-1338 | BdPV4 | EnaPV7 | PvlaPV3 | FcPV1 | AnPV1 | CaRNAV1 | AflPV1 | BdV1 |
|---|---|---|---|---|---|---|---|---|---|---|---|---|---|---|---|---|---|---|---|---|---|---|---|---|---|---|---|---|---|---|---|---|---|---|---|---|---|---|---|---|---|---|---|---|---|---|---|---|---|---|---|---|---|---|---|---|---|---|---|---|---|---|---|---|---|---|
| HPBV | | 10 | 11 | 11 | 11 | 11 | 9 | 10 | 10 | 10 | 9 | 10 | 10 | 9 | 11 | 9 | 9 | 10 | 11 | 11 | 10 | 10 | 10 | 10 | 10 | 11 | 10 | 10 | 12 | 12 | 13 | 13 | 12 | 12 | 11 | 12 | 13 | 12 | 13 | 11 | 14 | 13 | 13 | 13 | 13 | 13 | 13 | 14 | 14 | 13 | 13 | 15 | 15 | 14 | 14 | 14 | 15 | 14 | 14 | 15 | 16 | 15 | 13 | 14 | 14 | 14 | 13 |
| HetRV2 | 10 | | 65 | 38 | 38 | 38 | 35 | 33 | 34 | 35 | 34 | 34 | 33 | 34 | 35 | 32 | 33 | 32 | 21 | 19 | 21 | 20 | 21 | 20 | 20 | 22 | 23 | 20 | 14 | 13 | 14 | 15 | 15 | 16 | 15 | 14 | 14 | 14 | 13 | 14 | 13 | 12 | 13 | 13 | 14 | 13 | 13 | 13 | 13 | 13 | 14 | 14 | 15 | 14 | 15 | 15 | 14 | 16 | 16 | 15 | 15 | 15 | 14 | 15 | 15 | 15 | 14 |
| HetRV7 | 11 | 65 | | 39 | 38 | 38 | 36 | 31 | 33 | 34 | 33 | 33 | 34 | 33 | 36 | 36 | 34 | 34 | 22 | 21 | 20 | 20 | 20 | 22 | 21 | 23 | 23 | 20 | 14 | 15 | 15 | 17 | 14 | 16 | 15 | 15 | 14 | 14 | 14 | 14 | 15 | 13 | 13 | 14 | 13 | 13 | 14 | 14 | 14 | 14 | 14 | 15 | 16 | 15 | 15 | 16 | 15 | 16 | 16 | 15 | 16 | 16 | 15 | 15 | 15 | 15 | 14 |
| AhV | 11 | 38 | 39 | | 60 | 59 | 37 | 32 | 33 | 35 | 34 | 33 | 34 | 34 | 37 | 40 | 37 | 37 | 20 | 21 | 21 | 21 | 21 | 20 | 21 | 19 | 21 | 22 | 13 | 14 | 13 | 14 | 17 | 15 | 15 | 14 | 13 | 12 | 13 | 15 | 15 | 14 | 15 | 15 | 16 | 15 | 14 | 15 | 15 | 15 | 15 | 14 | 14 | 15 | 15 | 16 | 14 | 15 | 15 | 15 | 15 | 14 | 14 | 15 | 15 | 15 | 13 |
| CpPV | 11 | 38 | 38 | 60 | | 95 | 36 | 32 | 33 | 34 | 35 | 35 | 34 | 34 | 37 | 37 | 35 | 37 | 21 | 19 | 19 | 20 | 20 | 19 | 21 | 19 | 20 | 21 | 16 | 14 | 15 | 13 | 16 | 16 | 14 | 14 | 14 | 14 | 15 | 15 | 16 | 14 | 14 | 14 | 16 | 14 | 14 | 15 | 15 | 15 | 16 | 14 | 14 | 15 | 15 | 15 | 15 | 15 | 15 | 14 | 14 | 15 | 13 | 16 | 15 | 15 | 14 |
| CrV1 | 11 | 38 | 38 | 59 | 95 | | 36 | 33 | 33 | 35 | 35 | 35 | 35 | 35 | 37 | 37 | 35 | 38 | 21 | 19 | 19 | 19 | 19 | 19 | 20 | 18 | 20 | 21 | 16 | 15 | 14 | 13 | 16 | 16 | 15 | 14 | 14 | 14 | 14 | 15 | 16 | 14 | 14 | 15 | 16 | 14 | 14 | 15 | 15 | 15 | 16 | 15 | 14 | 15 | 15 | 15 | 16 | 16 | 16 | 15 | 15 | 15 | 13 | 16 | 15 | 15 | 14 |
| RsV | 9 | 35 | 36 | 37 | 36 | 36 | | 46 | 47 | 47 | 49 | 49 | 48 | 49 | 47 | 47 | 50 | 50 | 22 | 19 | 19 | 20 | 20 | 20 | 21 | 20 | 21 | 20 | 13 | 13 | 13 | 14 | 14 | 14 | 14 | 15 | 15 | 14 | 14 | 14 | 14 | 15 | 16 | 16 | 16 | 17 | 16 | 16 | 16 | 17 | 16 | 16 | 15 | 15 | 15 | 16 | 16 | 17 | 17 | 16 | 16 | 16 | 14 | 15 | 14 | 15 | 14 |
| CCV | 10 | 33 | 31 | 32 | 32 | 33 | 46 | | 77 | 76 | 78 | 78 | 77 | 76 | 53 | 48 | 49 | 49 | 22 | 18 | 20 | 18 | 19 | 18 | 19 | 20 | 21 | 17 | 14 | 13 | 13 | 15 | 14 | 14 | 15 | 16 | 15 | 15 | 14 | 14 | 14 | 14 | 15 | 15 | 15 | 14 | 16 | 16 | 16 | 14 | 15 | 15 | 14 | 15 | 15 | 16 | 15 | 16 | 16 | 15 | 15 | 15 | 15 | 15 | 15 | 14 | 13 |
| CCCV2 | 10 | 34 | 33 | 33 | 33 | 33 | 47 | 77 | | 77 | 79 | 81 | 80 | 79 | 52 | 49 | 48 | 47 | 23 | 20 | 22 | 22 | 22 | 21 | 22 | 20 | 23 | 19 | 15 | 12 | 13 | 14 | 14 | 15 | 16 | 17 | 14 | 14 | 14 | 16 | 16 | 15 | 15 | 15 | 15 | 15 | 16 | 15 | 15 | 15 | 15 | 16 | 15 | 16 | 15 | 16 | 16 | 16 | 16 | 15 | 15 | 15 | 16 | 17 | 17 | 16 | 15 |
| PmV1 | 10 | 35 | 34 | 35 | 34 | 35 | 47 | 76 | 77 | | 79 | 77 | 78 | 78 | 52 | 50 | 50 | 50 | 23 | 18 | 21 | 21 | 20 | 20 | 21 | 21 | 22 | 19 | 16 | 13 | 13 | 14 | 15 | 15 | 16 | 16 | 15 | 14 | 13 | 15 | 14 | 15 | 15 | 15 | 15 | 14 | 15 | 15 | 15 | 15 | 15 | 15 | 13 | 15 | 14 | 16 | 14 | 15 | 15 | 14 | 14 | 15 | 15 | 16 | 16 | 15 | 14 |
| DCV2 | 9 | 34 | 33 | 34 | 35 | 35 | 49 | 78 | 79 | 79 | | 83 | 81 | 80 | 53 | 51 | 50 | 50 | 23 | 20 | 21 | 20 | 20 | 19 | 20 | 21 | 23 | 19 | 15 | 13 | 13 | 14 | 14 | 14 | 15 | 15 | 15 | 15 | 14 | 16 | 16 | 15 | 15 | 15 | 15 | 16 | 15 | 15 | 15 | 15 | 15 | 16 | 15 | 16 | 17 | 17 | 16 | 17 | 17 | 16 | 16 | 17 | 16 | 17 | 16 | 15 | 15 |
| HTCV2 | 10 | 34 | 33 | 33 | 35 | 35 | 49 | 78 | 81 | 77 | 83 | | 87 | 85 | 53 | 51 | 50 | 51 | 23 | 20 | 22 | 20 | 20 | 20 | 21 | 21 | 23 | 19 | 15 | 13 | 13 | 14 | 14 | 15 | 16 | 16 | 15 | 14 | 15 | 16 | 16 | 15 | 15 | 15 | 15 | 15 | 15 | 15 | 15 | 16 | 15 | 15 | 14 | 15 | 15 | 15 | 15 | 16 | 16 | 15 | 15 | 15 | 15 | 16 | 16 | 15 | 15 |
| RCCV2 | 10 | 33 | 34 | 34 | 34 | 35 | 48 | 77 | 80 | 78 | 81 | 87 | | 91 | 53 | 51 | 50 | 49 | 22 | 20 | 22 | 21 | 21 | 20 | 21 | 22 | 23 | 20 | 15 | 12 | 13 | 14 | 13 | 15 | 16 | 16 | 16 | 15 | 14 | 17 | 16 | 15 | 15 | 15 | 14 | 15 | 16 | 15 | 16 | 15 | 15 | 16 | 14 | 16 | 15 | 16 | 16 | 16 | 16 | 15 | 15 | 16 | 16 | 17 | 17 | 15 | 16 |
| WCCV2 | 9 | 34 | 33 | 34 | 34 | 35 | 49 | 76 | 79 | 78 | 80 | 85 | 91 | | 52 | 50 | 50 | 50 | 23 | 20 | 22 | 21 | 20 | 20 | 21 | 21 | 23 | 20 | 16 | 13 | 13 | 14 | 14 | 15 | 15 | 16 | 16 | 14 | 15 | 17 | 16 | 15 | 16 | 15 | 15 | 16 | 16 | 16 | 16 | 16 | 16 | 15 | 14 | 15 | 15 | 16 | 15 | 16 | 16 | 15 | 15 | 15 | 15 | 16 | 17 | 15 | 15 |
| RnV1 | 11 | 35 | 36 | 37 | 37 | 37 | 47 | 53 | 52 | 52 | 53 | 53 | 53 | 52 | | 60 | 56 | 52 | 21 | 19 | 19 | 18 | 18 | 19 | 19 | 21 | 21 | 20 | 13 | 13 | 13 | 14 | 12 | 15 | 14 | 14 | 14 | 14 | 12 | 14 | 14 | 14 | 14 | 14 | 15 | 14 | 14 | 14 | 15 | 14 | 14 | 15 | 15 | 14 | 14 | 15 | 15 | 16 | 16 | 15 | 15 | 15 | 14 | 15 | 15 | 14 | 13 |
| PoV1 | 9 | 32 | 36 | 40 | 37 | 37 | 47 | 48 | 49 | 50 | 51 | 51 | 51 | 50 | 60 | | 64 | 63 | 23 | 20 | 21 | 20 | 20 | 20 | 20 | 21 | 23 | 21 | 14 | 13 | 13 | 15 | 14 | 15 | 14 | 16 | 15 | 14 | 13 | 16 | 16 | 16 | 14 | 14 | 15 | 14 | 15 | 14 | 15 | 14 | 15 | 15 | 14 | 15 | 14 | 16 | 14 | 15 | 15 | 15 | 15 | 15 | 15 | 15 | 14 | 13 | 13 |
| FpV1 | 9 | 33 | 34 | 37 | 35 | 35 | 50 | 49 | 48 | 50 | 50 | 50 | 50 | 50 | 56 | 64 | | 71 | 23 | 19 | 20 | 18 | 18 | 18 | 18 | 20 | 20 | 20 | 13 | 11 | 12 | 13 | 13 | 15 | 15 | 16 | 14 | 14 | 14 | 15 | 14 | 15 | 14 | 14 | 14 | 13 | 14 | 14 | 14 | 13 | 14 | 14 | 14 | 13 | 13 | 14 | 13 | 14 | 14 | 14 | 14 | 14 | 14 | 14 | 14 | 14 | 13 |
| HetRV8 | 10 | 32 | 34 | 37 | 37 | 38 | 50 | 49 | 47 | 50 | 50 | 51 | 49 | 50 | 52 | 63 | 71 | | 21 | 18 | 20 | 19 | 19 | 19 | 20 | 20 | 20 | 20 | 12 | 12 | 12 | 14 | 14 | 15 | 15 | 15 | 15 | 15 | 15 | 15 | 15 | 14 | 15 | 15 | 15 | 15 | 15 | 16 | 16 | 14 | 15 | 16 | 14 | 14 | 15 | 16 | 15 | 16 | 16 | 15 | 15 | 15 | 14 | 15 | 15 | 15 | 14 |
| RnPV2 | 11 | 21 | 22 | 20 | 21 | 21 | 22 | 22 | 23 | 23 | 23 | 23 | 22 | 23 | 21 | 23 | 23 | 21 | | 28 | 29 | 28 | 29 | 29 | 29 | 29 | 33 | 31 | 14 | 15 | 15 | 15 | 17 | 18 | 18 | 16 | 15 | 15 | 17 | 18 | 17 | 16 | 16 | 17 | 17 | 17 | 17 | 16 | 16 | 16 | 17 | 19 | 17 | 19 | 18 | 18 | 17 | 18 | 18 | 17 | 18 | 19 | 18 | 18 | 18 | 18 | 17 |
| HetPV1 | 11 | 19 | 21 | 21 | 19 | 19 | 19 | 18 | 20 | 18 | 20 | 20 | 20 | 20 | 19 | 20 | 19 | 18 | 28 | | 51 | 56 | 54 | 55 | 53 | 30 | 34 | 34 | 16 | 13 | 14 | 14 | 16 | 16 | 16 | 15 | 15 | 15 | 16 | 16 | 16 | 16 | 16 | 17 | 16 | 16 | 18 | 18 | 17 | 15 | 16 | 17 | 15 | 16 | 15 | 15 | 14 | 15 | 15 | 16 | 15 | 15 | 14 | 15 | 15 | 15 | 15 |
| CCRSaPV | 10 | 21 | 20 | 21 | 19 | 19 | 19 | 20 | 22 | 21 | 21 | 22 | 22 | 22 | 19 | 21 | 20 | 20 | 29 | 51 | | 58 | 59 | 60 | 59 | 31 | 35 | 33 | 14 | 14 | 14 | 13 | 16 | 15 | 15 | 14 | 15 | 15 | 15 | 16 | 16 | 15 | 15 | 15 | 15 | 15 | 15 | 15 | 14 | 16 | 15 | 15 | 15 | 15 | 16 | 17 | 16 | 16 | 16 | 16 | 16 | 16 | 15 | 15 | 16 | 16 | 17 |
| BCV1 | 10 | 20 | 20 | 21 | 20 | 19 | 20 | 18 | 22 | 21 | 20 | 20 | 21 | 21 | 18 | 20 | 18 | 19 | 28 | 56 | 58 | | 84 | 80 | 82 | 31 | 36 | 35 | 14 | 12 | 13 | 14 | 15 | 14 | 15 | 15 | 13 | 14 | 14 | 15 | 16 | 14 | 16 | 15 | 15 | 15 | 16 | 16 | 15 | 15 | 16 | 17 | 15 | 14 | 15 | 15 | 15 | 15 | 15 | 16 | 15 | 15 | 14 | 15 | 15 | 16 | 16 |
| CrCV | 10 | 21 | 20 | 21 | 20 | 19 | 20 | 19 | 22 | 20 | 20 | 20 | 21 | 20 | 18 | 20 | 18 | 19 | 29 | 54 | 59 | 84 | | 80 | 83 | 32 | 36 | 37 | 13 | 14 | 13 | 13 | 15 | 14 | 15 | 14 | 13 | 14 | 14 | 15 | 16 | 13 | 14 | 15 | 15 | 14 | 15 | 15 | 15 | 14 | 16 | 16 | 15 | 15 | 15 | 15 | 15 | 15 | 15 | 16 | 16 | 16 | 14 | 15 | 15 | 16 | 16 |
| VCV | 10 | 20 | 22 | 20 | 19 | 19 | 20 | 18 | 21 | 20 | 19 | 20 | 20 | 20 | 19 | 20 | 18 | 19 | 29 | 55 | 60 | 80 | 80 | | 83 | 31 | 35 | 36 | 13 | 14 | 14 | 13 | 15 | 14 | 15 | 14 | 12 | 13 | 14 | 14 | 16 | 14 | 14 | 14 | 14 | 14 | 14 | 15 | 14 | 14 | 15 | 17 | 15 | 15 | 16 | 15 | 15 | 15 | 15 | 16 | 15 | 15 | 14 | 14 | 15 | 15 | 16 |
| WCCV1 | 10 | 20 | 21 | 21 | 21 | 20 | 21 | 19 | 22 | 21 | 20 | 21 | 21 | 21 | 19 | 20 | 18 | 20 | 29 | 53 | 59 | 82 | 83 | 83 | | 32 | 36 | 37 | 14 | 14 | 14 | 13 | 16 | 14 | 15 | 14 | 13 | 13 | 14 | 15 | 16 | 14 | 14 | 15 | 15 | 14 | 15 | 15 | 15 | 15 | 16 | 17 | 16 | 16 | 16 | 16 | 16 | 16 | 16 | 17 | 17 | 17 | 14 | 15 | 15 | 17 | 17 |
| HetRV3 | 11 | 22 | 23 | 19 | 19 | 18 | 20 | 20 | 20 | 21 | 21 | 21 | 22 | 21 | 21 | 21 | 20 | 20 | 29 | 30 | 31 | 31 | 32 | 31 | 32 | | 31 | 34 | 14 | 12 | 15 | 13 | 17 | 15 | 15 | 14 | 14 | 13 | 12 | 15 | 15 | 12 | 13 | 14 | 14 | 13 | 14 | 14 | 13 | 13 | 14 | 15 | 13 | 15 | 14 | 16 | 13 | 13 | 13 | 13 | 14 | 14 | 14 | 14 | 15 | 15 | 15 |
| CpCV1 | 10 | 23 | 23 | 21 | 20 | 20 | 21 | 21 | 23 | 22 | 23 | 23 | 23 | 23 | 21 | 23 | 20 | 20 | 33 | 34 | 35 | 36 | 36 | 35 | 36 | 31 | | 45 | 14 | 14 | 13 | 14 | 16 | 16 | 17 | 18 | 17 | 15 | 18 | 19 | 19 | 18 | 16 | 17 | 16 | 17 | 17 | 16 | 16 | 17 | 17 | 18 | 17 | 17 | 17 | 17 | 17 | 17 | 17 | 17 | 17 | 17 | 17 | 18 | 17 | 17 | 18 |
| FvBV | 10 | 20 | 20 | 22 | 21 | 21 | 20 | 17 | 19 | 19 | 19 | 19 | 20 | 20 | 20 | 21 | 20 | 20 | 31 | 34 | 33 | 35 | 37 | 36 | 37 | 34 | 45 | | 15 | 14 | 14 | 14 | 15 | 15 | 16 | 15 | 15 | 15 | 15 | 16 | 16 | 14 | 15 | 15 | 15 | 17 | 16 | 15 | 15 | 15 | 15 | 18 | 15 | 17 | 17 | 17 | 17 | 17 | 17 | 17 | 18 | 18 | 17 | 15 | 16 | 16 | 16 |
| PCV1 | 12 | 14 | 14 | 13 | 16 | 16 | 13 | 14 | 15 | 16 | 15 | 15 | 15 | 16 | 13 | 14 | 13 | 12 | 14 | 16 | 14 | 14 | 13 | 13 | 14 | 14 | 14 | 15 | | 53 | 51 | 16 | 17 | 17 | 17 | 17 | 17 | 17 | 21 | 18 | 18 | 19 | 17 | 18 | 19 | 19 | 19 | 19 | 19 | 18 | 18 | 21 | 18 | 18 | 19 | 19 | 19 | 19 | 19 | 21 | 19 | 19 | 19 | 17 | 19 | 19 | 18 |
| BCV2 | 12 | 13 | 15 | 14 | 14 | 15 | 13 | 13 | 12 | 13 | 13 | 13 | 12 | 13 | 13 | 13 | 11 | 12 | 15 | 13 | 14 | 12 | 14 | 14 | 14 | 12 | 14 | 14 | 53 | | 55 | 16 | 15 | 16 | 15 | 18 | 18 | 18 | 18 | 16 | 16 | 17 | 17 | 18 | 19 | 18 | 18 | 19 | 19 | 18 | 17 | 17 | 17 | 18 | 17 | 18 | 18 | 18 | 18 | 18 | 17 | 18 | 18 | 16 | 17 | 18 | 16 |
| PCV2 | 13 | 14 | 15 | 13 | 15 | 14 | 13 | 13 | 13 | 13 | 13 | 13 | 13 | 13 | 13 | 13 | 12 | 12 | 15 | 14 | 14 | 13 | 13 | 14 | 14 | 15 | 13 | 14 | 51 | 55 | | 18 | 17 | 17 | 17 | 17 | 18 | 18 | 20 | 18 | 18 | 17 | 17 | 17 | 17 | 17 | 17 | 17 | 17 | 17 | 17 | 17 | 16 | 18 | 18 | 18 | 18 | 18 | 18 | 18 | 18 | 18 | 18 | 17 | 17 | 18 | 18 |
| CSpV1 | 13 | 15 | 17 | 14 | 13 | 13 | 14 | 15 | 14 | 14 | 14 | 14 | 14 | 14 | 14 | 15 | 13 | 14 | 15 | 14 | 13 | 14 | 13 | 13 | 13 | 13 | 14 | 14 | 16 | 16 | 18 | | 15 | 16 | 16 | 17 | 17 | 16 | 15 | 15 | 14 | 18 | 17 | 17 | 18 | 17 | 18 | 18 | 18 | 20 | 19 | 16 | 18 | 18 | 17 | 17 | 16 | 18 | 18 | 18 | 18 | 18 | 16 | 16 | 16 | 16 | 15 |
| RsdRV5 | 12 | 15 | 14 | 17 | 16 | 16 | 14 | 14 | 14 | 15 | 14 | 14 | 13 | 14 | 12 | 14 | 13 | 14 | 17 | 16 | 16 | 15 | 15 | 15 | 16 | 17 | 16 | 15 | 17 | 15 | 17 | 15 | | 40 | 40 | 16 | 17 | 16 | 15 | 15 | 16 | 17 | 16 | 16 | 17 | 16 | 16 | 16 | 16 | 16 | 16 | 19 | 18 | 19 | 20 | 19 | 19 | 19 | 19 | 18 | 18 | 19 | 18 | 19 | 18 | 19 | 18 |
| BbPV3 | 12 | 16 | 16 | 15 | 16 | 16 | 14 | 14 | 15 | 15 | 14 | 15 | 14 | 15 | 15 | 15 | 15 | 15 | 18 | 16 | 15 | 14 | 14 | 14 | 14 | 15 | 16 | 15 | 17 | 16 | 17 | 16 | 40 | | 56 | 17 | 17 | 17 | 18 | 17 | 17 | 17 | 18 | 17 | 17 | 18 | 18 | 18 | 18 | 18 | 18 | 21 | 19 | 19 | 19 | 19 | 18 | 20 | 20 | 19 | 20 | 20 | 16 | 19 | 19 | 18 | 18 |
| CePV1 | 11 | 15 | 15 | 15 | 14 | 15 | 14 | 15 | 16 | 16 | 15 | 16 | 16 | 15 | 14 | 14 | 15 | 15 | 18 | 16 | 15 | 15 | 15 | 15 | 15 | 15 | 17 | 16 | 17 | 15 | 17 | 16 | 40 | 56 | | 16 | 19 | 16 | 18 | 18 | 17 | 18 | 18 | 19 | 20 | 18 | 18 | 19 | 19 | 18 | 18 | 21 | 21 | 20 | 21 | 21 | 21 | 21 | 21 | 20 | 20 | 20 | 19 | 19 | 20 | 19 | 19 |
| FePV1 | 12 | 14 | 15 | 14 | 14 | 14 | 15 | 16 | 17 | 16 | 15 | 16 | 16 | 16 | 14 | 16 | 16 | 15 | 16 | 15 | 14 | 15 | 14 | 14 | 14 | 14 | 18 | 15 | 17 | 18 | 17 | 17 | 16 | 17 | 16 | | 65 | 64 | 19 | 21 | 20 | 22 | 24 | 24 | 23 | 23 | 24 | 24 | 24 | 23 | 22 | 21 | 19 | 20 | 19 | 20 | 19 | 20 | 20 | 20 | 20 | 19 | 21 | 21 | 20 | 22 | 20 |
| AaPV1 | 13 | 14 | 14 | 13 | 14 | 14 | 15 | 15 | 14 | 15 | 15 | 15 | 16 | 16 | 14 | 15 | 14 | 15 | 15 | 15 | 15 | 13 | 13 | 12 | 13 | 14 | 17 | 15 | 17 | 18 | 18 | 17 | 17 | 17 | 19 | 65 | | 73 | 19 | 19 | 19 | 23 | 23 | 23 | 22 | 23 | 24 | 23 | 23 | 24 | 22 | 21 | 21 | 20 | 21 | 21 | 20 | 20 | 21 | 19 | 21 | 20 | 20 | 21 | 21 | 22 | 21 |
| BdPV1_ | 12 | 14 | 14 | 12 | 14 | 14 | 14 | 15 | 14 | 14 | 15 | 14 | 15 | 14 | 15 | 14 | 14 | 15 | 15 | 15 | 15 | 14 | 14 | 13 | 13 | 13 | 15 | 15 | 17 | 18 | 18 | 16 | 16 | 17 | 16 | 64 | 73 | | 19 | 19 | 18 | 22 | 23 | 23 | 23 | 24 | 23 | 23 | 23 | 23 | 22 | 21 | 21 | 21 | 21 | 20 | 20 | 22 | 22 | 19 | 20 | 20 | 20 | 20 | 20 | 21 | 20 |
| PnV1 | 13 | 13 | 14 | 13 | 15 | 14 | 14 | 14 | 14 | 13 | 14 | 15 | 14 | 15 | 12 | 13 | 14 | 15 | 17 | 16 | 15 | 14 | 14 | 14 | 14 | 12 | 18 | 15 | 21 | 18 | 20 | 15 | 15 | 18 | 18 | 19 | 19 | 19 | | 48 | 50 | 26 | 24 | 25 | 23 | 25 | 25 | 25 | 25 | 25 | 24 | 22 | 21 | 19 | 21 | 20 | 21 | 21 | 21 | 21 | 19 | 20 | 21 | 22 | 21 | 23 | 22 |
| MoPV1 | 11 | 14 | 14 | 15 | 15 | 15 | 14 | 15 | 16 | 15 | 16 | 16 | 17 | 17 | 14 | 16 | 15 | 15 | 18 | 16 | 16 | 15 | 15 | 14 | 15 | 15 | 19 | 16 | 18 | 16 | 18 | 15 | 15 | 17 | 18 | 21 | 19 | 19 | 48 | | 63 | 24 | 25 | 25 | 25 | 25 | 26 | 26 | 26 | 24 | 24 | 21 | 22 | 21 | 21 | 21 | 21 | 22 | 22 | 22 | 20 | 21 | 22 | 23 | 22 | 25 | 23 |
| PsV-F | 14 | 13 | 15 | 15 | 16 | 16 | 14 | 14 | 16 | 14 | 16 | 16 | 16 | 16 | 14 | 16 | 14 | 15 | 17 | 16 | 16 | 16 | 16 | 16 | 16 | 15 | 19 | 16 | 18 | 16 | 18 | 14 | 16 | 17 | 17 | 20 | 19 | 18 | 50 | 63 | | 25 | 24 | 25 | 24 | 25 | 25 | 25 | 24 | 24 | 25 | 21 | 21 | 20 | 20 | 19 | 21 | 20 | 20 | 21 | 19 | 20 | 21 | 23 | 23 | 24 | 23 |
| FsV1 | 13 | 12 | 13 | 14 | 14 | 14 | 15 | 14 | 15 | 15 | 15 | 15 | 15 | 15 | 14 | 16 | 15 | 14 | 16 | 16 | 15 | 14 | 13 | 14 | 14 | 12 | 18 | 14 | 19 | 17 | 17 | 18 | 17 | 17 | 18 | 22 | 23 | 22 | 26 | 24 | 25 | | 58 | 57 | 60 | 59 | 59 | 58 | 58 | 59 | 60 | 28 | 28 | 29 | 29 | 28 | 28 | 28 | 29 | 27 | 27 | 27 | 27 | 27 | 28 | 29 | 28 |
| OPV1 | 13 | 13 | 13 | 15 | 14 | 14 | 16 | 14 | 15 | 15 | 15 | 15 | 15 | 16 | 14 | 14 | 14 | 15 | 16 | 16 | 15 | 16 | 14 | 14 | 14 | 13 | 16 | 15 | 17 | 17 | 17 | 17 | 16 | 18 | 18 | 24 | 23 | 23 | 24 | 25 | 24 | 58 | | 72 | 72 | 71 | 73 | 74 | 73 | 65 | 67 | 28 | 27 | 29 | 29 | 29 | 28 | 28 | 28 | 28 | 29 | 28 | 28 | 30 | 30 | 30 | 29 |
| DdV1 | 13 | 13 | 14 | 15 | 14 | 15 | 16 | 15 | 15 | 15 | 15 | 15 | 15 | 15 | 14 | 14 | 14 | 15 | 17 | 17 | 15 | 15 | 15 | 14 | 15 | 14 | 17 | 15 | 18 | 18 | 17 | 17 | 16 | 17 | 19 | 24 | 23 | 23 | 25 | 25 | 25 | 57 | 72 | | 89 | 79 | 81 | 81 | 80 | 64 | 70 | 29 | 28 | 29 | 29 | 28 | 29 | 29 | 29 | 28 | 27 | 28 | 27 | 29 | 29 | 29 | 28 |
| DdV2 | 13 | 14 | 13 | 16 | 16 | 16 | 16 | 15 | 15 | 15 | 15 | 15 | 14 | 15 | 15 | 15 | 14 | 15 | 17 | 16 | 15 | 15 | 15 | 14 | 15 | 14 | 16 | 15 | 19 | 19 | 17 | 18 | 17 | 17 | 20 | 23 | 22 | 23 | 23 | 25 | 24 | 60 | 72 | 89 | | 80 | 82 | 81 | 80 | 64 | 68 | 29 | 27 | 28 | 28 | 27 | 28 | 28 | 28 | 27 | 26 | 26 | 25 | 29 | 28 | 28 | 27 |
| UvPV | 13 | 13 | 13 | 15 | 14 | 14 | 17 | 14 | 15 | 14 | 16 | 15 | 15 | 16 | 14 | 14 | 13 | 15 | 17 | 16 | 15 | 15 | 14 | 14 | 14 | 13 | 17 | 17 | 19 | 18 | 17 | 17 | 16 | 18 | 18 | 23 | 23 | 24 | 25 | 25 | 25 | 59 | 71 | 79 | 80 | | 85 | 83 | 83 | 66 | 70 | 30 | 29 | 29 | 30 | 29 | 30 | 30 | 30 | 29 | 29 | 29 | 28 | 30 | 29 | 31 | 29 |
| CcPV1-1338 | 13 | 13 | 14 | 14 | 14 | 14 | 16 | 16 | 16 | 15 | 15 | 15 | 16 | 16 | 14 | 15 | 14 | 15 | 17 | 18 | 15 | 16 | 15 | 14 | 15 | 14 | 17 | 16 | 19 | 18 | 17 | 18 | 16 | 18 | 18 | 24 | 24 | 23 | 25 | 26 | 25 | 59 | 73 | 81 | 82 | 85 | | 91 | 91 | 65 | 71 | 31 | 29 | 29 | 29 | 29 | 29 | 30 | 30 | 28 | 28 | 28 | 28 | 31 | 30 | 30 | 29 |
| CcPV1-Cb3-2 | 14 | 13 | 14 | 15 | 15 | 15 | 16 | 16 | 15 | 15 | 15 | 15 | 15 | 16 | 14 | 14 | 14 | 16 | 16 | 18 | 15 | 16 | 15 | 15 | 15 | 14 | 16 | 15 | 19 | 19 | 17 | 18 | 16 | 18 | 19 | 24 | 23 | 23 | 25 | 26 | 25 | 58 | 74 | 81 | 81 | 83 | 91 | | 98 | 65 | 69 | 29 | 28 | 27 | 28 | 27 | 28 | 29 | 28 | 27 | 27 | 27 | 27 | 30 | 29 | 29 | 28 |
| EnaPV3 | 14 | 13 | 14 | 15 | 15 | 15 | 16 | 16 | 15 | 15 | 15 | 15 | 16 | 16 | 15 | 15 | 14 | 16 | 16 | 17 | 14 | 15 | 15 | 14 | 15 | 13 | 16 | 15 | 19 | 19 | 17 | 18 | 16 | 18 | 19 | 24 | 23 | 23 | 25 | 26 | 24 | 58 | 73 | 80 | 80 | 83 | 91 | 98 | | 64 | 69 | 29 | 28 | 28 | 28 | 27 | 28 | 29 | 29 | 27 | 27 | 27 | 27 | 29 | 29 | 29 | 28 |
| AoV | 13 | 13 | 14 | 15 | 15 | 15 | 17 | 14 | 15 | 15 | 15 | 16 | 15 | 16 | 14 | 14 | 13 | 14 | 16 | 15 | 16 | 15 | 14 | 14 | 15 | 13 | 17 | 15 | 18 | 18 | 17 | 20 | 16 | 18 | 18 | 23 | 24 | 23 | 25 | 24 | 24 | 59 | 65 | 64 | 64 | 66 | 65 | 65 | 64 | | 72 | 28 | 28 | 30 | 29 | 29 | 28 | 29 | 29 | 27 | 29 | 29 | 27 | 29 | 28 | 30 | 30 |
| PsV-S | 13 | 14 | 14 | 15 | 16 | 16 | 16 | 15 | 15 | 15 | 15 | 15 | 15 | 16 | 14 | 15 | 14 | 15 | 17 | 16 | 15 | 16 | 16 | 15 | 16 | 14 | 17 | 15 | 18 | 17 | 17 | 19 | 16 | 18 | 18 | 22 | 22 | 22 | 24 | 24 | 25 | 60 | 67 | 70 | 68 | 70 | 71 | 69 | 69 | 72 | | 29 | 29 | 29 | 30 | 29 | 30 | 31 | 30 | 28 | 29 | 29 | 27 | 30 | 29 | 29 | 28 |
| UvPV2 | 15 | 14 | 15 | 14 | 14 | 15 | 16 | 15 | 16 | 15 | 16 | 15 | 16 | 15 | 15 | 15 | 14 | 16 | 19 | 17 | 15 | 17 | 16 | 17 | 17 | 15 | 18 | 18 | 21 | 17 | 17 | 16 | 19 | 21 | 21 | 21 | 21 | 21 | 22 | 21 | 21 | 28 | 28 | 29 | 29 | 30 | 31 | 29 | 29 | 28 | 29 | | 60 | 65 | 63 | 63 | 65 | 64 | 64 | 65 | 65 | 65 | 47 | 48 | 48 | 50 | 47 |
| UvMV | 15 | 15 | 16 | 14 | 14 | 14 | 15 | 14 | 15 | 13 | 15 | 14 | 14 | 14 | 15 | 14 | 14 | 14 | 17 | 15 | 15 | 15 | 15 | 15 | 16 | 13 | 17 | 15 | 18 | 17 | 16 | 18 | 18 | 19 | 21 | 19 | 21 | 21 | 21 | 22 | 21 | 28 | 27 | 28 | 27 | 29 | 29 | 28 | 28 | 28 | 29 | 60 | | 66 | 65 | 66 | 71 | 64 | 64 | 65 | 66 | 67 | 51 | 53 | 51 | 51 | 50 |
| ThPV2 | 14 | 14 | 15 | 15 | 15 | 15 | 15 | 15 | 16 | 15 | 16 | 15 | 16 | 15 | 14 | 15 | 13 | 14 | 19 | 16 | 15 | 14 | 15 | 15 | 16 | 15 | 17 | 17 | 18 | 18 | 18 | 18 | 19 | 19 | 20 | 20 | 20 | 21 | 19 | 21 | 20 | 29 | 29 | 29 | 28 | 29 | 29 | 27 | 28 | 30 | 29 | 65 | 66 | | 76 | 77 | 71 | 73 | 73 | 71 | 74 | 74 | 50 | 49 | 51 | 50 | 49 |
| CaPV1 | 14 | 15 | 15 | 15 | 15 | 15 | 15 | 15 | 15 | 14 | 17 | 15 | 15 | 15 | 14 | 14 | 13 | 15 | 18 | 15 | 16 | 15 | 15 | 16 | 16 | 14 | 17 | 17 | 19 | 17 | 18 | 17 | 20 | 19 | 21 | 19 | 21 | 21 | 21 | 21 | 20 | 29 | 29 | 29 | 28 | 30 | 29 | 28 | 28 | 29 | 30 | 63 | 65 | 76 | | 89 | 71 | 73 | 73 | 69 | 73 | 73 | 48 | 51 | 51 | 51 | 51 |
| CgPV1 | 14 | 15 | 16 | 16 | 15 | 15 | 16 | 16 | 16 | 16 | 17 | 15 | 16 | 16 | 15 | 16 | 14 | 16 | 18 | 15 | 17 | 15 | 15 | 15 | 16 | 16 | 17 | 17 | 19 | 18 | 18 | 17 | 19 | 19 | 21 | 20 | 21 | 20 | 20 | 21 | 19 | 28 | 29 | 28 | 27 | 29 | 29 | 27 | 27 | 29 | 29 | 63 | 66 | 77 | 89 | | 71 | 73 | 72 | 70 | 72 | 72 | 50 | 51 | 52 | 51 | 51 |
| MpPV1 | 15 | 14 | 15 | 14 | 15 | 16 | 16 | 15 | 16 | 14 | 16 | 15 | 16 | 15 | 15 | 14 | 13 | 15 | 17 | 14 | 16 | 15 | 15 | 15 | 16 | 13 | 17 | 17 | 19 | 18 | 18 | 16 | 19 | 18 | 21 | 19 | 20 | 20 | 21 | 21 | 21 | 28 | 28 | 29 | 28 | 30 | 29 | 28 | 28 | 28 | 30 | 65 | 71 | 71 | 71 | 71 | | 74 | 74 | 73 | 74 | 74 | 50 | 52 | 52 | 54 | 52 |
| PvlaPV4 | 14 | 16 | 16 | 15 | 15 | 16 | 17 | 16 | 16 | 15 | 17 | 16 | 16 | 16 | 16 | 15 | 14 | 16 | 18 | 15 | 16 | 15 | 15 | 15 | 16 | 13 | 17 | 17 | 19 | 18 | 18 | 18 | 19 | 20 | 21 | 20 | 20 | 22 | 21 | 22 | 20 | 28 | 28 | 29 | 28 | 30 | 30 | 29 | 29 | 29 | 31 | 64 | 64 | 73 | 73 | 73 | 74 | | 99 | 75 | 76 | 77 | 51 | 51 | 51 | 51 | 51 |
| CcPV2-1338 | 14 | 16 | 16 | 15 | 15 | 16 | 17 | 16 | 16 | 15 | 17 | 16 | 16 | 16 | 16 | 15 | 14 | 16 | 18 | 15 | 16 | 15 | 15 | 15 | 16 | 13 | 17 | 17 | 19 | 18 | 18 | 18 | 19 | 20 | 21 | 20 | 21 | 22 | 21 | 22 | 20 | 29 | 28 | 29 | 28 | 30 | 30 | 28 | 29 | 29 | 30 | 64 | 64 | 73 | 73 | 72 | 74 | 99 | | 75 | 77 | 77 | 51 | 51 | 51 | 51 | 51 |
| BdPV4 | 15 | 15 | 15 | 15 | 14 | 15 | 16 | 15 | 15 | 14 | 16 | 15 | 15 | 15 | 15 | 15 | 14 | 15 | 17 | 16 | 16 | 16 | 16 | 16 | 17 | 13 | 17 | 17 | 21 | 18 | 18 | 18 | 18 | 19 | 20 | 20 | 20 | 19 | 19 | 21 | 22 | 27 | 28 | 28 | 27 | 29 | 28 | 27 | 27 | 27 | 28 | 65 | 65 | 71 | 69 | 70 | 73 | 75 | 75 | | 80 | 80 | 50 | 52 | 51 | 53 | 52 |
| EnaPV7 | 16 | 15 | 16 | 15 | 14 | 15 | 16 | 15 | 15 | 14 | 16 | 15 | 15 | 15 | 15 | 15 | 14 | 15 | 18 | 15 | 16 | 15 | 16 | 15 | 17 | 14 | 17 | 18 | 19 | 17 | 18 | 18 | 18 | 20 | 20 | 20 | 21 | 20 | 19 | 20 | 19 | 27 | 29 | 27 | 26 | 29 | 28 | 27 | 27 | 29 | 29 | 65 | 66 | 74 | 73 | 72 | 74 | 76 | 77 | 80 | | 97 | 50 | 50 | 51 | 51 | 50 |
| PvlaPV3 | 15 | 15 | 16 | 14 | 15 | 15 | 16 | 15 | 15 | 15 | 17 | 15 | 16 | 15 | 15 | 15 | 14 | 15 | 19 | 15 | 16 | 15 | 16 | 15 | 17 | 14 | 17 | 18 | 19 | 18 | 18 | 18 | 19 | 20 | 20 | 19 | 20 | 20 | 20 | 21 | 20 | 27 | 28 | 28 | 26 | 29 | 28 | 27 | 27 | 29 | 29 | 65 | 67 | 74 | 73 | 72 | 74 | 77 | 77 | 80 | 97 | | 51 | 50 | 51 | 51 | 50 |
| FcPV1 | 13 | 14 | 15 | 14 | 13 | 13 | 14 | 15 | 16 | 15 | 16 | 15 | 16 | 15 | 14 | 15 | 14 | 14 | 18 | 14 | 15 | 14 | 14 | 14 | 14 | 14 | 17 | 17 | 19 | 18 | 17 | 16 | 18 | 16 | 19 | 21 | 20 | 20 | 21 | 22 | 21 | 27 | 28 | 27 | 25 | 28 | 28 | 27 | 27 | 27 | 27 | 47 | 51 | 50 | 48 | 50 | 50 | 51 | 51 | 50 | 50 | 51 | | 57 | 56 | 55 | 56 |
| AnPV1 | 14 | 15 | 15 | 15 | 16 | 16 | 15 | 15 | 17 | 16 | 17 | 16 | 17 | 16 | 15 | 15 | 14 | 15 | 18 | 15 | 15 | 15 | 15 | 14 | 15 | 14 | 18 | 15 | 17 | 16 | 17 | 16 | 19 | 19 | 19 | 21 | 21 | 20 | 22 | 23 | 23 | 27 | 30 | 29 | 29 | 30 | 31 | 30 | 29 | 29 | 30 | 48 | 53 | 49 | 51 | 51 | 52 | 51 | 51 | 52 | 50 | 50 | 57 | | 73 | 76 | 77 |
| CaRNAV1 | 14 | 15 | 15 | 15 | 15 | 15 | 14 | 15 | 17 | 16 | 16 | 16 | 17 | 17 | 15 | 14 | 14 | 15 | 18 | 15 | 16 | 15 | 15 | 15 | 15 | 15 | 17 | 16 | 19 | 17 | 18 | 16 | 18 | 19 | 20 | 20 | 21 | 20 | 21 | 22 | 23 | 28 | 30 | 29 | 28 | 29 | 30 | 29 | 29 | 28 | 29 | 48 | 51 | 51 | 51 | 52 | 52 | 51 | 51 | 51 | 51 | 51 | 56 | 73 | | 76 | 77 |
| AflPV1 | 14 | 15 | 15 | 15 | 15 | 15 | 15 | 14 | 16 | 15 | 15 | 15 | 15 | 15 | 14 | 13 | 14 | 15 | 18 | 15 | 16 | 16 | 16 | 15 | 17 | 15 | 17 | 16 | 19 | 18 | 18 | 16 | 19 | 18 | 19 | 22 | 22 | 21 | 23 | 25 | 24 | 29 | 30 | 29 | 28 | 31 | 30 | 29 | 29 | 30 | 29 | 50 | 51 | 50 | 51 | 51 | 54 | 51 | 51 | 53 | 51 | 51 | 55 | 76 | 76 | | 83 |
| BdV1 | 13 | 14 | 14 | 13 | 14 | 14 | 14 | 13 | 15 | 14 | 15 | 15 | 16 | 15 | 13 | 13 | 13 | 14 | 17 | 15 | 17 | 16 | 16 | 16 | 17 | 15 | 18 | 16 | 18 | 16 | 18 | 15 | 18 | 18 | 19 | 20 | 21 | 20 | 22 | 23 | 23 | 28 | 29 | 28 | 27 | 29 | 29 | 28 | 28 | 30 | 28 | 47 | 50 | 49 | 51 | 51 | 52 | 51 | 51 | 52 | 50 | 50 | 56 | 77 | 77 | 83 | |
