## Supplementary material for "New viruses of *Cladosporium* sp. expand considerably the taxonomic structure of *Gammapartitivirus* genus": Fig. S

A

| Extraction | Illumina HTS | Reads | Bases QC >30 (%) | Contigs (>100bp) | N50 | Candidates | Validated by RT-PCR |
|---|---|---|---|---|---|---|---|
| Total-RNA | mRNA | 646,123,786 | 94.23 | 625,876 | 296 | 15 | 1 |
| Total-RNA | sRNA | 107,320,035 | 90.23 | - | - | - | - |
| VANA | mRNA | 18,053 | 89.39 | 287 | 234 | 4 | 4 |

B

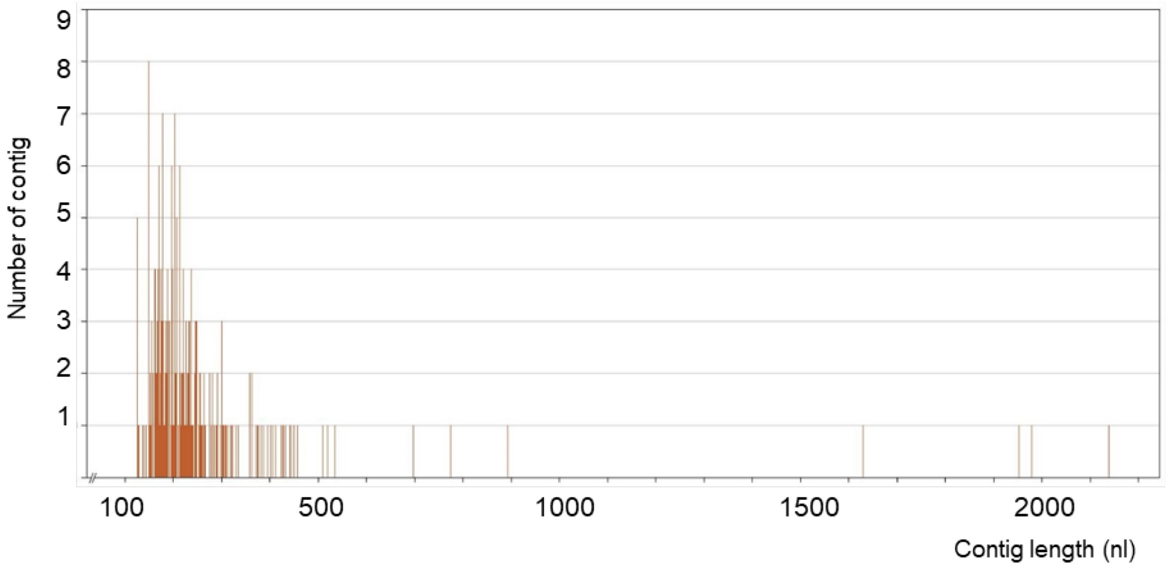


**Fig. S1: Sequencing data information.** A] Output data from HTS analysis of the fungal community of Leytron (Total-RNA) and the *C. cladospirioides* AGS-1338 from the Agroscope collection (VANA). Reads were assembled with geneious assembler, and blastn on a local database with a selection of mycovirus sequences. RT-PCR was performed with primer designed on the selected candidates (primers of validated sequences in the Table S1). B] Barplot of the contigs assembled with Geneious assembler of strain C. cladospirioides AGS-1338 from the *de novo* assembly of the VANA RNA extraction HTS reads.

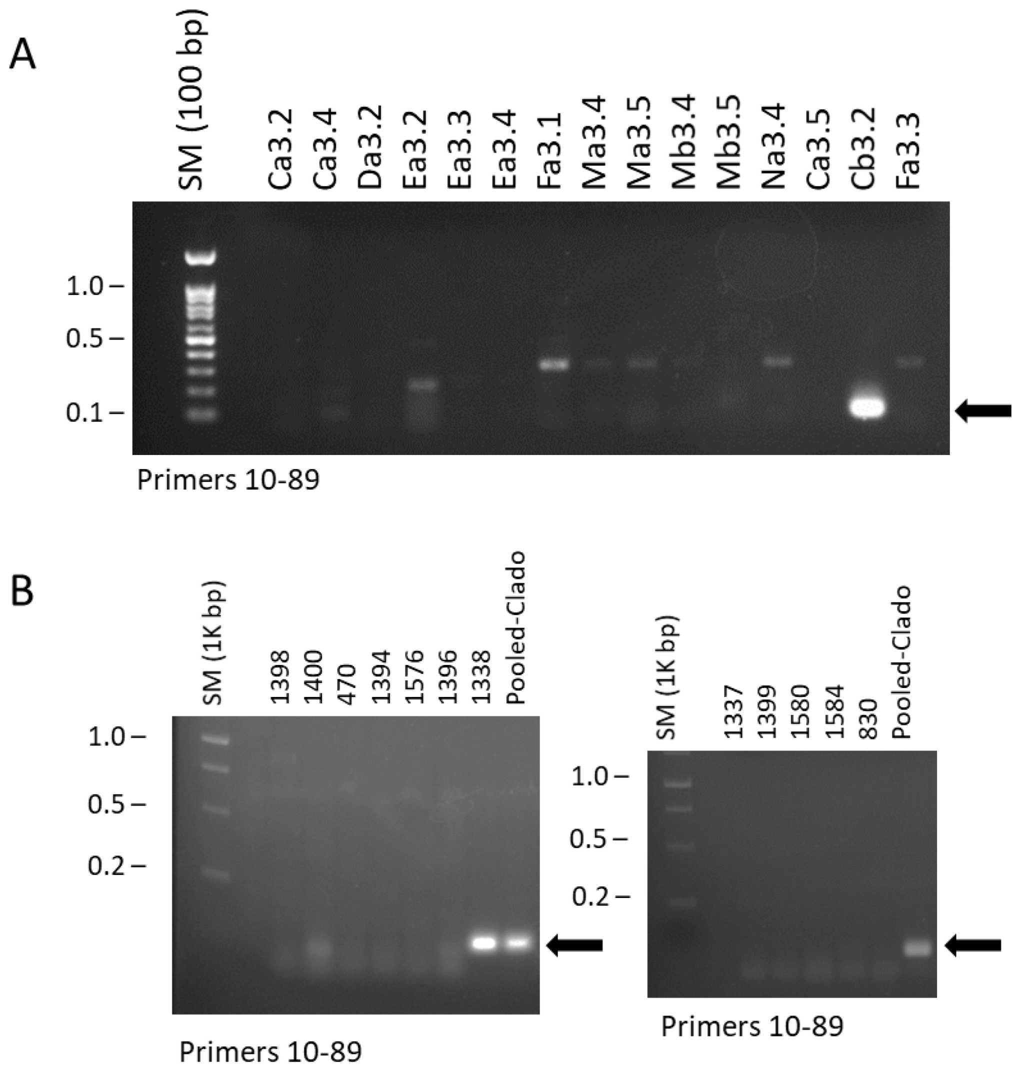


**Fig. S2 : Detection of the CcPV1**. RT-PCR with primer designed on the RdRp of CcPV1. The primer sequences are in the Supplementary Table S1, and expected length is highlighted with a white arrow. The size marker (SM) is a DNA ladder of 100 bp (A) and 1Kbp (B). A] Screen for the presence of CcPV1 in RNA extracts from isolates of the collection of Leytron. B] (both panels) Screen for the presence of CcPV1 in the pool of all *Cladosporium sp.* RNA extracts sent for Illumina sequencing (pooled –C. sp) of the collection of Leytron and in individual RNA extracts from Cladosporium sp. from the Agroscope collection (B).
